# Discovering nuclear localization signal universe through a novel deep learning model with interpretable attention units

**DOI:** 10.1101/2024.08.10.606103

**Authors:** Yi-Fan Li, Xiaoyong Pan, Hong-Bin Shen

## Abstract

Nuclear localization signals (NLSs) are essential peptide fragments within proteins that play a decisive role in guiding proteins into the cell nucleus. Determining the existence and precise locations of NLSs experimentally is time-consuming and complicated, resulting in a scarcity of experimentally validated NLS fragments. Consequently, annotated NLS datasets are relatively limited, presenting challenges for data-driven approaches. In this study, we propose an innovative interpretable approach, NLSExplorer, which leverages large-scale protein language models to capture crucial biological information with a novel attention-based deep network for NLS identification. By enhancing the knowledge retrieved from protein language models with a novel attention to key area module, NLSExplorer achieves superior predictive performance compared to existing methods on two NLS benchmark datasets. Additionally, NLSExplorer is able to detect various kinds of segments highly correlated with nuclear transport, such as nuclear export signals. We employ NLSExplorer to investigate potential NLSs and other domains that are important for nuclear transport in nucleus-localized proteins in the Swiss-Prot database. Furthermore, the comprehensive pattern analysis for all these segments uncovers a potential NLS space and internal relationship of important nuclear transport segments for 416 species. This study not only introduces a powerful tool for predicting and exploring NLS space, but also offers a versatile network that is powerful for detecting characteristic domains and motifs of NLSs.

## Introduction

The subcellular localization^1^ of proteins refers to the positions of proteins within the various organelles of a biological cell. It is a crucial factor in determining the functions of proteins. Nuclear localization signal (NLS)^2^ sequences are amino acid fragments that are responsible for the cellular nucleus targeting and transporting. NLSs are discovered in conjunction with the corresponding transport proteins, ultimately allowing transport proteins to enter the interior of the cell nucleus by interacting with nucleoporins^3^. The exploration and comprehensive understanding of NLSs can facilitate the revelation of the principles underlying the complex cellular processes^4-6^. In recent years, the association of NLS sequences and their variations with the disease occurrence has been increasingly investigated, promoting the development of related clinical drugs^7^. For instance, the role of NLSs in viral infection and nuclear transport has contributed to the development of therapeutic vaccines^8^ and novel antiviral drugs^9, 10^. However, experimental detection of NLS signals imposes high demands on experimental techniques and costs^11-14^, resulting in a limited understanding of NLSs at present.

To speed up the understanding of NLSs, various computational methods have been progressively developed (refer to Supplementary Table 1). These methods primarily fall into three categories: Template-matching, machine learning, and electronic mutation-based methods. These computational methods are fundamentally dependent on the experimental dataset size, but the availability of NLS data remains limited. To date, there are a total of 14,416 experimentally validated sequences located within the cell nucleus in the Swiss-Prot (2024_04)^15^ database. Of them, the number of sequences with experimentally validated NLS is only 298. Thus, the performance of existing methods for predicting NLS is limited. It is mainly due to the following factors: (1) Methods are easily over-fitted and lack sufficient understanding of NLS patterns when the training dataset contains only a small number of NLSs. This significantly limits data-driven approaches like machine learning, resulting in poor generalization performance. (2) Template-matching approaches and electronic mutation-based methods utilize experimentally validated NLSs to form templates or as mutation objects, causing them to seek and discover typical sequence segments defined by experts. However, due to the significant diversity in NLSs, it is challenging to identify novel patterns and unseen NLSs.

Protein language models have opened a new way to address learning problems with limited task-specific protein sequences. The key of a protein language model is to train on a large scale of unlabeled proteins, enabling it to gain understanding for different kinds of language like human language, codex language, or biology language. This training process is centered on the exploration of the statistic rules implied in amino acids and capturing semantic correlations^16^ among diverse subjects. These features captured by language models contain rich information on proteins about evolution, function, and structure. By leveraging this characteristic, protein language models provide new and unique ways to explore biological knowledge^17^. Computational predictors combined with large language models have demonstrated superior performance to traditional methods in different fields of biology like structure prediction^18^, missense variant effect prediction^19^, peptide-detection^20^, subcellular localization prediction^21^. Despite the abundance of biological knowledge integrated into the language model to achieve excellent performance, for specific downstream tasks, it is still difficult to know which specific information has been utilized in these tasks. In contrary, biologists prefer to seek such information^22^ for biological interpretation.

To address the aforementioned limitations and challenges, we propose an interpretable approach NLSExplorer, which consists of two main parts, for NLS prediction and pattern discovery. The first part is the prediction system NLSExplorer-prediction equipped with an explorer module called Attention to Key Area (A2KA). It identifies crucial areas by extracting biological information from the embedding space of large language models. We train A2KA model to distinguish the nuclear localizations of proteins in Swiss-Prot using representations generated by ESM1b-650M. This process enables the model with the ability of large-scale detection for nuclear transport segments within entire sequences, especially NLSs. This alleviates the limitations on discovering new types of NLSs due to the scarcity of experimental NLSs data. Subsequently, a recommendation system is cascaded to provide more precise screening for NLSs utilizing the key segments generated by A2KA. Such a mode of “explore + screen” enables NLSExplorer-prediction to achieve a high accuracy in NLS prediction on two benchmark datasets with over 10% F1 score improvement. Furthermore, we utilize NLSExplorer-prediction to thoroughly detect potential NLSs in Swiss-Prot database. By using these potential segments, NLSExplorer has the ability for NLS prediction and similar NLS segment search simultaneously. The retrieved similar NLS segments exhibit conservation on sequence and structure with predicted NLSs, providing an overview of the NLSs cross-species evolution. Based on these potential segments, we build an online interactive map called the NLS Candidate Library with global and local search function, shedding light on the potential universe of NLSs.

The second part NLSExplorer-SCNLS of NLSExplorer involves the Search and Collect NLS (SCNLS) algorithm for post-analysis of recommended segments. This algorithm is primarily designed to detect NLSs patterns, demonstrating capabilities for mining discontinuous NLS patterns. We use NLSExplorer-SCNLS to analyze the patterns of recommended nuclear transport segments by A2KA. Leveraging the analysis results, we develop another map called the Nuclear Transport Pattern Map to showcase frequent potential patterns of segments that play key role in nuclear transport and their characteristics, like diversity distribution. In addition, we explore the nuclear transport segments from the perspective of species, unveil a potential multi-species relationship for 416 species in Swiss-Prot. This study provides a novel approach with the exploration capability originating from the attention mechanism and the protein language model, exploring the boundaries of the NLS prediction.

## Results

### Overview of NLSExplorer

The core idea behind NLSExplorer is to fully leverage knowledge extracted from a protein language model to enhance NLS identification. As illustrated in Figure 1-a, we draw an analogy between an intuitive knowledge extraction scenario and our large language model-based approach. The aim is to understand the knowledge about protein sequences to use by an expert while dealing with different tasks. Instead of asking the expert to explicitly explain their decision process, we employ a direct observational approach. The task for expert is set to distinguishing nuclear proteins from a large set of proteins. By monitoring where the expert’s attention lingers the longest with a recorder and pattern analysis system, we can track their focus during the prediction process. After the expert has made prediction across a diverse set of proteins, which may include various structures, motifs, and domains associated with protein functions, the areas of the focus are highlighted, revealing the preferences of this expert.

**Figure 1.**
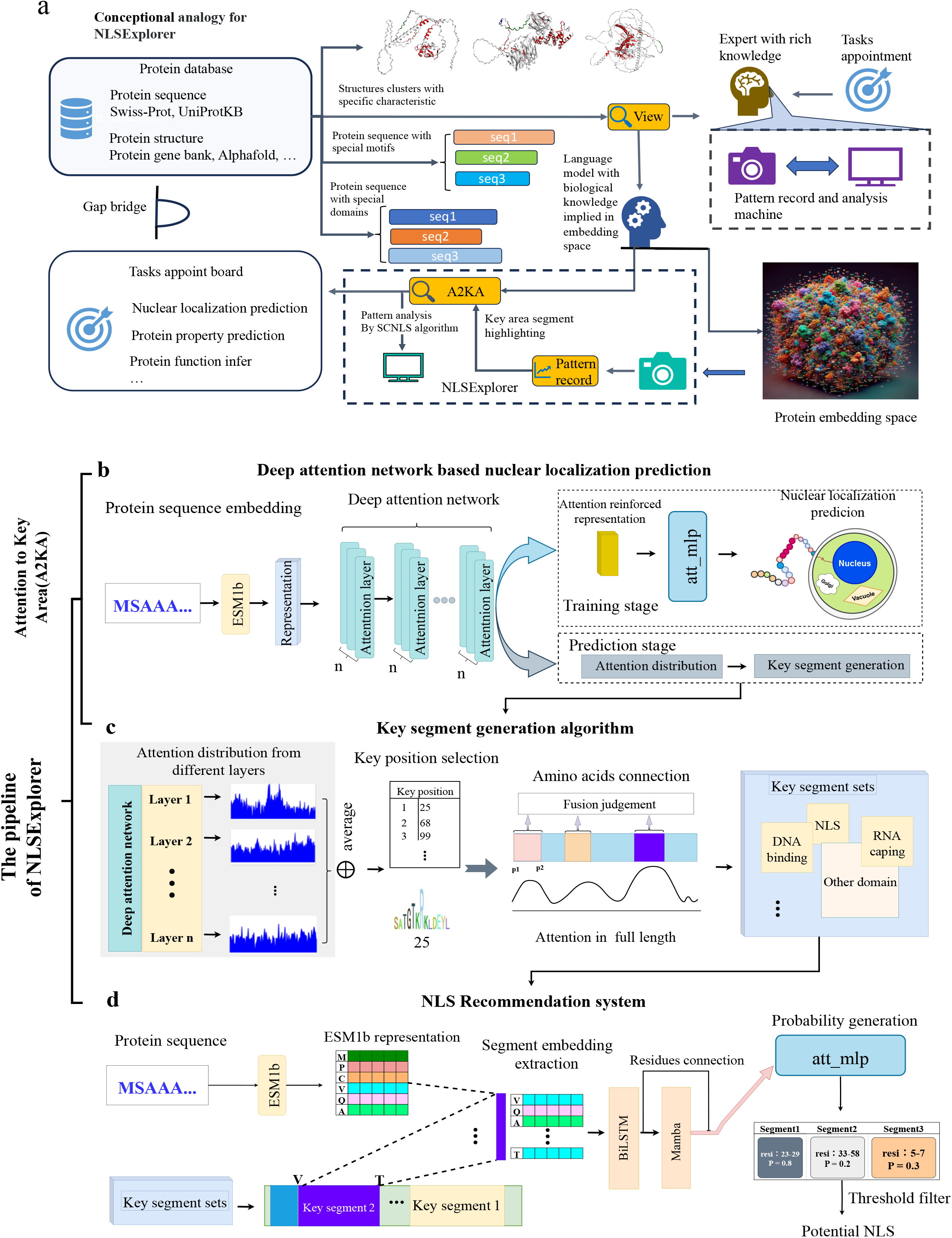
The schematic overview of NLSExplorer. **(a)** The conceptual analogy behind the motivation for NLSExplorer. **(b)** NLSExplorer-prediction makes predictions for NLS based on an attention mechanism with direct interpretability. It consists of A2KA (Attention to Key Area) and an NLS recommendation system. DAN (Deep Attention Network) is the first component of A2KA, which primarily produces an attention distribution to evaluate the relative importance of each site as a potential NLS. **(c)** The key segment generation module is the second component of A2KA; it utilizes the attention from DAN to connect residues and form key segment sets for NLS candidates. **(d)** The NLS recommendation system is cascaded to A2KA and further screens actual NLSs from the key segment sets.

These preferences help the expert filter out irrelevant data, focusing on the protein segments that contain the most critical information for prediction. Additionally, they offer deep insights into the nuclear localization knowledge embedded in the expert’s decision-making. In our language model-based approach, training NLSExplorer on the entire set of nuclear protein sequences from Swiss-Prot effectively creates an ‘expert’ in nuclear localization prediction. When identifying nuclear proteins, the A2KA component of NLSExplorer records patterns in the protein regions where attention is focused on. The segments in these regions correspond to specific motifs that influence protein localization within the nucleus, which are highly likely to contain NLSs. This significantly narrows the search space and reduces the complexity of NLS detection. The well-filtered segments are then passed to a recommendation system for final prediction. Additionally, we employ NLSExplorer-analysis as a pattern analysis tool to investigate the characteristics of these segments across a broad range of nuclear proteins, further revealing key characteristic of nuclear transport regions.

### NLSExplorer-prediction module for NLS prediction

To be more specific, in NLSExplorer, NLS prediction is performed within a relatively comprehensive process that involves two crucial components (Figure 1-b and 1-c): a predictive framework A2KA with embedded interpretability for potential segment generation, and a recommendation system for NLS prediction. A2KA is developed based on attention mechanisms^23^, it is mainly used for two objectives. One is to reinforce the representation generated by protein language models according to the attention weights to support nucleus localization predictors. The other objective is to utilize attention maps generated by attention mechanisms to uncover amino acid segments in the key areas of the protein sequence that may significantly influence the prediction results. The second module of NLSExplorer-prediction is a recommendation system, the key segment sets generated from the first module A2KA are then fed into it to determine the probability of being an NLS, and the segments with high probability are recommended as potential NLSs.

### Attention to Key Area framework

#### Attention distribution generation with deep attention network

We choose ESM-1b-650M^24^ for protein representation generation. The ESM-1b-650M model has a basic Transformer structure with 33 layers, pre-trained on the UR50/S^25^ 2018_03 dataset through self-supervised learning. In the first module (Figure 1-b), the protein sequence is transformed into a representation matrix (*E*) through ESM-1B-650M. This matrix, *E*, is subsequently fed into a deep attention network (DAN) composed of basic attention units (BAU). DAN is trained using nuclear subcellular localization as the supervised label. It predicts subcellular localization primarily by using the attention map generated by BAU to focus on specific areas of the protein sequence. The attention distribution map obtained from the attention enhancement module serves as the input for the key segment generation module to extract segments from key areas.

#### Key segment generation for nuclear localization by a segment generation algorithm

The second module of A2KA is the key segment generation module (Figure 1-c), which extracts the attention distribution map from various layers of DAN and aggregates them into a total distribution. It selects amino acids that rank in the top exploration cofactor proportion in the attention matrix (i.e. an exploration cofactor of 0.3 means that for each prediction, 0.3\**l* residues with high attention weights than other residues are used to generate the key segments, where *l* represents the sequence length) and fuses them into amino acid segments within a defined distance *H*. This module autonomously generates candidate key segments, providing a preliminary candidate set of key regions for specific tasks.

### NLS Recommendation System

The second part is the recommendation system (Figure 1-d), which maximizes the utilization of preliminarily generated candidate set. The recommendation system module is connected to the key segment generation module (Figure 1-c), which generates important segments for prediction based on the attention distribution map. Next, these segments are fed into the recommendation system to predict their probabilities of being potential NLSs. Finally, these segments are ranked according to their probabilities.

### NLSExplorer-SCNLS module for continuous and discontinuous pattern mining

We employ the NLSExplorer model with the SCNLS algorithm (Methods) to analyze all protein sequences located within the nucleus with experimental evidence in the Swiss-Prot database. For a recommended set generated by NLSExplorer-prediction module, the SCNLS algorithm systematically detects recurrent occurrences of frequent continuous segments, highlighting segments that may play pivotal roles ^3^. The SCNLS algorithm can capture non-continuous segments for discovering discontinuous NLS patterns. For instance, a typical segment of NLS is R*RK^26^, where * can be any residue. Hence, capturing frequent discontinuous patterns becomes imperative in the context of potential NLS exploration.

### NLSExplorer Achieves Promising Performance in Identifying NLSs on Two Benchmark Datasets

We first evaluate NLSExplorer on the training dataset of INSP^27^ using 5-fold cross-validation to optimize the model hyperparameters with four metrics: precision, recall, F1 score, and Prediction-aPC (Methods), which is a metric used to evaluate the coverage between all the recommended segments and the actual NLSs in the prediction task. The hyperparameters that yield the best performance are selected for the final model training. We set a threshold for recommendation, where segments with a potential NLS probability higher than the threshold are recommended as NLSs. The model is evaluated over a range of 25 epochs, with detailed results in Extended Data Fig 1. It is observed that the model trained for 25 epochs achieves the highest F1 score, with stable performance varying with the threshold (Figure 2-a). Therefore, we choose a threshold of 0.7, where the model achieves the best F1 score and Prediction-aPC of 0.2 for 5-fold cross-validation. Finally, we train our model with 25 epochs for performance comparison with other predictors on the INSP test set.

**Figure 2.**
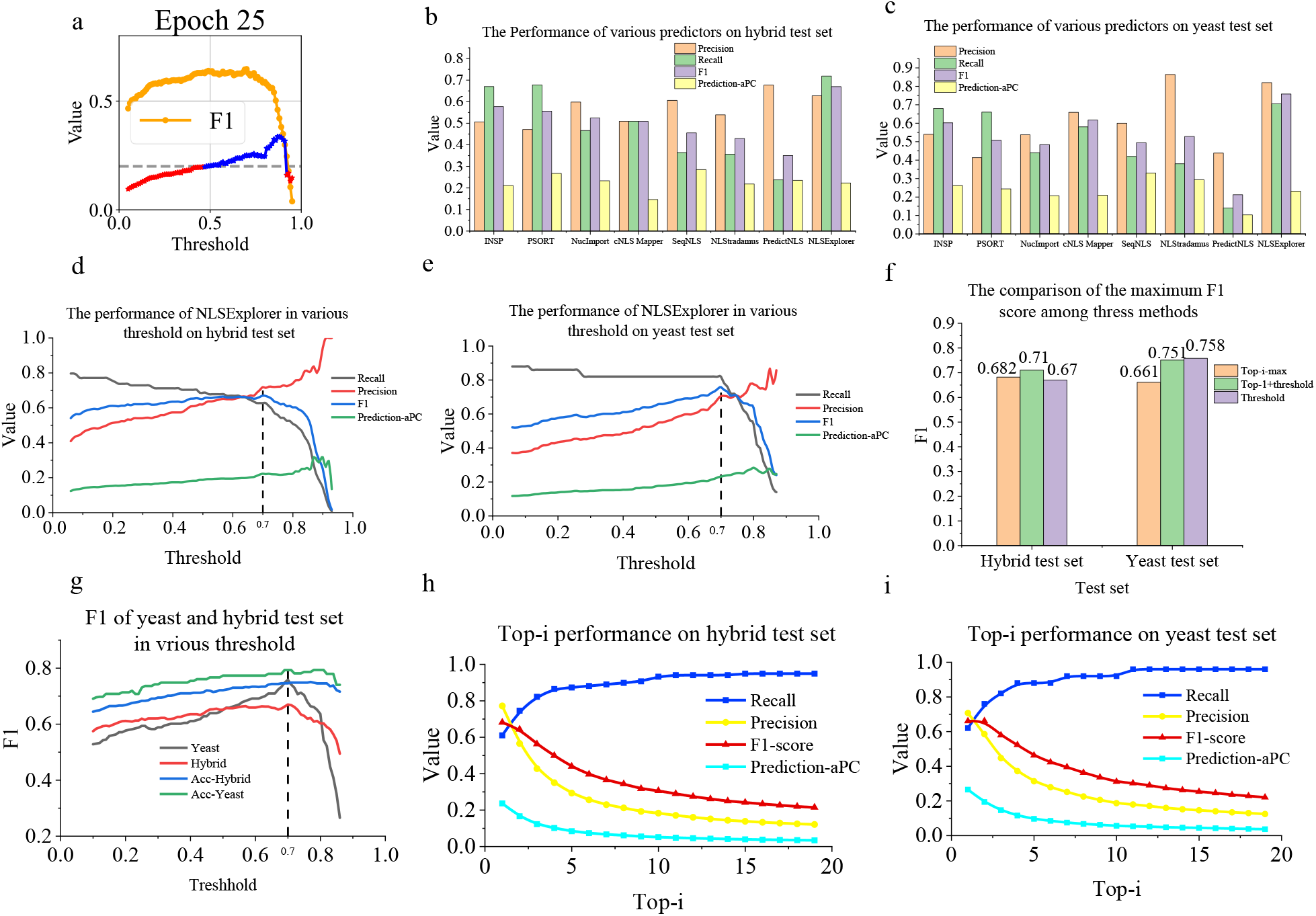
NLSExplorer achieves high performance on the INSP test dataset. **(a)** The Prediction-aPC and F1 score of NLSExplorer at the 25 epochs on the training dataset of INSP using 5-fold cross-validation. **(b)** The performance of various predictors on the hybrid test set. **(c)** The performance of various predictors on the yeast test set. **(d)** The performance of NLSExplorer with various thresholds on the hybrid test set. **(e)** The performance of NLSExplorer with various thresholds on the yeast test set. **(f)** The comparison of the maximum F1 score among three methods. **(g)** The F1 score of NLSExplorer-Accuracy and NLSExplorer-Lightning in the yeast and hybrid test set in various thresholds. The prefix “Acc” indicates the NLSExplorer-Accuracy model. **(h)** Top-i performance on the hybrid test set. **(i)** Top-i performance on the yeast test set.

Figures 2-b and 2-c show the performance of various predictors on the INSP hybrid and yeast test datasets. The hybrid dataset contains proteins with experimentally validated NLSs from multiple species, enabling the assessment of the model’s ability to generalize across species. NLSExplorer achieves the highest recall of 0.719 among all predictors, coupled with a precision of 0.627. NLSExplorer yields an F1 score of 0.670 with a 9% improvement compared to the previous best approach INSP^27^. Additionally, NLSExplorer achieves a Prediction-aPC of 0.22, placing it at a middle rank among all predictors. The yeast test set comprises proteins with experimentally validated NLSs from yeast species, which is a primary species for NLS study due to its abundant NLS data. NLSExplorer achieves a recall of 0.705, the highest among all predictors, and a precision of 0.82. NLSExplorer yields an F1 score of 0.758, marking a 14.1% improvement over the prior best approach cNLS Mapper^28^, which achieves an F1 score of 0.617. The results on these two datasets show that NLSExplorer achieves high recall with relatively high precision and Prediction-aPC.

To further investigate the impact of the classification threshold of the recommendation system on model performance, we apply different thresholds for NLS recommendation. Figure 2-d illustrates the performance across varying thresholds in the hybrid test set. As the threshold increases, both the Prediction-aPC and precision curves show an upward trend, while the recall decreases. The F1 score stabilizes and improves when the threshold is below 0.7 but sharply declines after 0.8, primarily due to a large drop in recall. Figure 2-e shows the performance across varying threshold in the yeast test set. The overall trend is similar to that of the hybrid test set. The F1 score shows a substantial decrease after the threshold exceeds 0.8. These results illustrate that NLSExplorer with a threshold of 0.7 achieves the best F1 score and falls within a range that avoids the sharp performance decline observed at the threshold of 0.8.

In addition to making accurate predictions for NLSs from protein sequences, another key goal of developing NLSExplorer is to create a model capable of exploring the NLS space. The INSP dataset contains proteins with at least one NLS, making it suitable for simulating scenarios where key NLSs are present in protein sequences. Using NLSExplorer, we employ four methods to facilitate NLS discovery, as shown in Figure 2-f.

1. Top-i: Recommends segments with the highest i probabilities of being NLS.
2. Top-i max: The best F1 score is achieved when selecting different i values.
3. Threshold: Recommends segments as NLSs if their probability exceeds the predefined threshold. In our comparisons, we use a threshold of 0.7 for classification.
4. Top-1 + Threshold: For each protein, it first recommends the segment with the highest NLS probability and then appends segments if their probability exceeds the threshold of 0.7.

In the hybrid test set, the Top-1 + Threshold method achieves the best F1 score with a 4% improvement compared to the Threshold method and a 3% improvement compared to the Top-i max method. In the yeast test set, the Threshold method achieves the best F1 score with a 10% improvement compared to the Top-i max method and a 0.7% improvement compared to the Top-1+Threshold method. The results demonstrate that the Threshold and Top-i max methods are suited for different scenarios. The mixed method, Top-1 + Threshold, shows relatively robust performance across both datasets. The detailed results of these methods are given in Extended Data Fig. 2.

The current recommendation system of NLSExplorer first extracts the embeddings of segments directly from the overall protein embedding, which avoids processing the entire sequence embedding and results in high computational efficiency. However, this method omits the surrounding information of the segment, leading to a loss of contextual information. We hypothesize that a balance between computational efficiency and the model performance can be achieved. Therefore, we conduct another experiment to validate this. Apart from the current structure, we design another structure for the recommendation system in NLSExplorer. We add a 3-layer Mamba network as the initial layer to first process the entire protein embedding, denoted as NLSExplorer-Accuracy. The model without the initial Mamba layer for comparison with other predictors in the earlier section is denoted as NLSExplorer-Lightning. We train NLSExplorer-Accuracy on the INSP training dataset and evaluate its performance on test datasets.

Figure 2-g shows a comparison of these two models on the INSP hybrid and yeast test datasets, demonstrating that NLSExplorer-Accuracy outperforms NLSExplorer-Lightning on both datasets, achieving an F1 score improvement of 8% and 3.5% with the 0.7 threshold, respectively. Furthermore, the overall F1 curve of the NLSExplorer-Accuracy model is more robust to threshold variations compared to the NLSExplorer-Lightning model. This implies that simply adding a Mamba layer to the initial input of the recommendation system produces a more distinct representation of the recommended segments by aggregating overall information, which benefits the NLS classification. However, although the NLSExplorer-Accuracy model yields better performance, it requires significantly more training steps, as well as longer computational time (see Extended Data Fig. 2). When the resources are sufficient, the NLSExplorer-Accuracy model is a good choice. Otherwise, the lightning model can be a suitable option. In addition, we test the performance of Top-1+Threshold for NLSExplorer-Accuracy model to evaluate the influence of adding the Top-1 recommendation into the threshold result for NLSExplorer-Accuracy, as shown in Extended Data Figure. 2.

Figures 2-h and 2-i show the Top-i results in the hybrid and yeast test sets. The model achieves a recall of approximately 0.9 when i is set to 5 on both datasets. Although recall continues to improve, the F1 score drops significantly as i increases. For exploration purpose, we need a high recall to capture potential NLSs and new types of NLSs. Furthermore, the threshold method shows a high accuracy in NLS prediction and provides an assessment of the likelihood that a segment is an NLS. Therefore, we ultimately use Top-5 + Threshold with the NLSExplorer-Accuracy mode of NLSExplorer to explore the NLS space and construct the NLS candidate maps.

### A2KA in NLSExplorer Uncovers Important Areas in Protein Sequences

In this section, we demonstrate that the protein subcellular location labels for training A2KA enable its attention mechanism to detect not only NLSs, but also other important motifs or domains. As shown in Figure 3-a, A2KA exhibits a hierarchical detection capability across protein regions. It prioritizes areas most associated with nuclear localization, such as NLSs, followed by segments partially related to nuclear localization. Even though, the interference from unrelated area can be introduced while increasing the exploration scope. We also show that the NLSs recommended by NLSExplorer share sequence and structural similarities, suggesting a common function. This recommendation feature enhances prediction results by highlighting potential NLS segments with similar properties during the prediction process, providing an overall reference to explore the potential NLS space.

**Figure 3.**
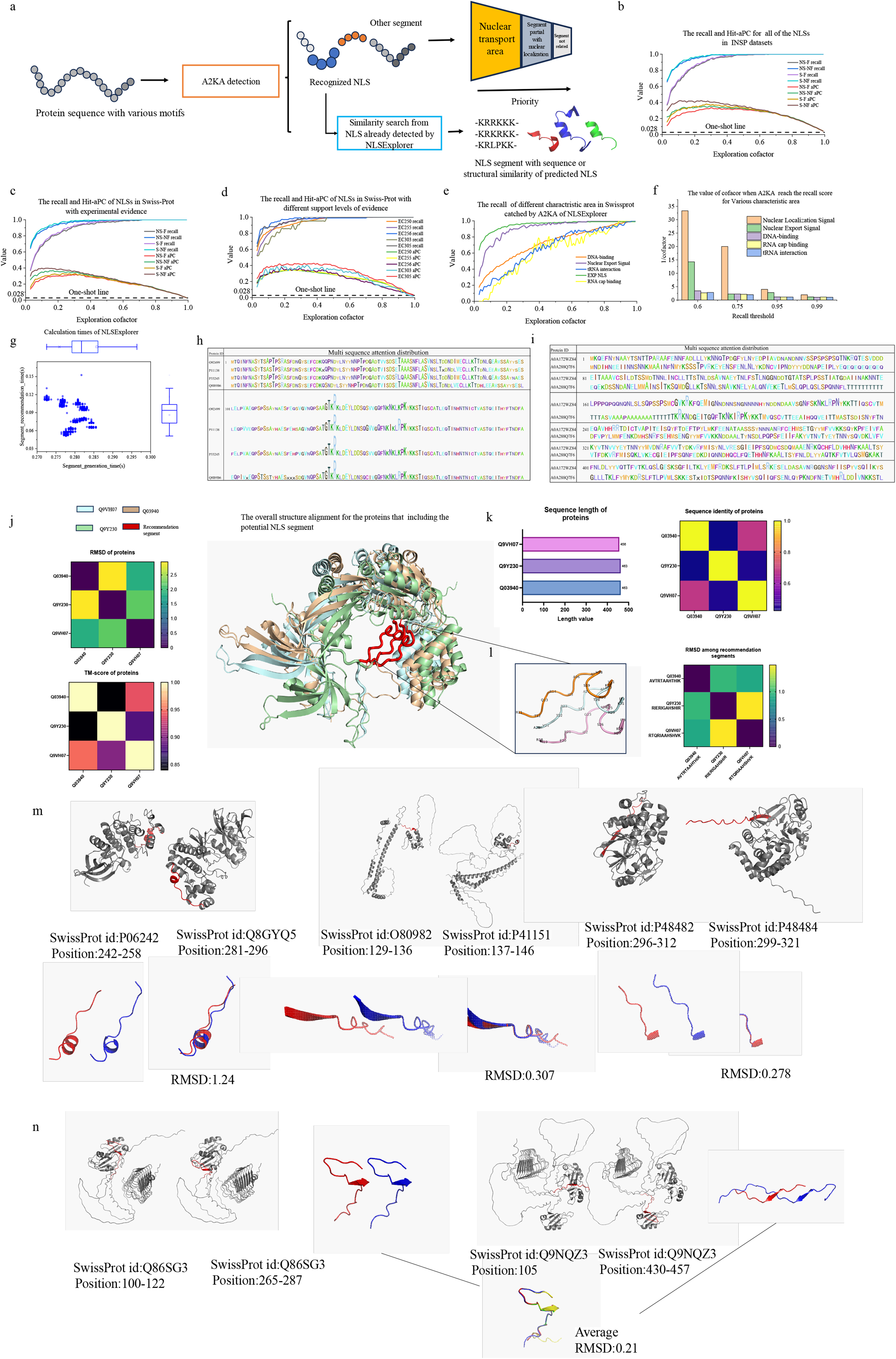
Overall performance of NLSExplorer on various tasks. **(a)** The recommendation characteristics of NLSExplorer. **(b)** The recall and Hit-aPC of A2KA with different parameters in the INSP dataset. The one-shot line indicates the Hit-aPC by predicting the entire protein sequence as an NLS. **(c)** The recall and Hit-aPC of A2KA for NLSs in the Swiss-Prot dataset with experimental evidence. **(d)** The recall and Hit-aPC of A2KA for NLSs with different support levels of evidence. **(e)** The recall of A2KA with different characteristic areas in Swiss-Prot. **(f)** The value of exploration cofactor when A2KA reaches different recalls for various characteristic areas. **(g)** The computational time of NLSExplorer. **(h)** Multiple sequence attentions for protein P11138 with the E-value of 0 and High-scoring Segment Pair (HSP) identity is over 90%. **(i)** Multiple sequence attentions for MSA of P11138 with E-value of 0 but HSP identity is less than 90%. **(j)** The structure alignment for three proteins which are detected by NLSExplorer to be the potential NLS segment. RMSD (Root Mean Square Deviation) and TM-score are displayed at the bottom. **(k)** The sequence length and identity of the three proteins, the sequence identity is calculated by BLAST. **(l)** The aligned recommended NLS segment shows a minimum and maximum RMSD of 0.89 and 1.49, respectively, indicating high similarity. **(m)** The alignment displays the recommended NLSs alongside their nearest neighbor in the embedding space of the recommendation system. **(n)** The DAZ (Deleted in Azoospermia) protein contains two recommended NLSs with highly similar structures.

### A2KA discovers different characteristic regions on protein sequences

We apply A2KA on all the NLSs data with various support levels in Swiss-Prot and conduct predictions on proteins with several selected motifs to evaluate its exploratory abilities. These selected motifs include NES (Nuclear Export Signals), DNA-binding, tRNA interaction, and RNA cap binding. They encompass not only regions that are highly correlated with protein transfer in and out nucleus, such as NES, but also regions involved in biological processes like tRNA interactions that do not fully occur in the nucleus. We collect these data from Swiss-Prot, which contains the term “ECO:0000269” as clear evidence of experimental validation. We use the recall and Hit-aPC (Methods) as metrics to test the influence of two important parameters of A2KA on NLS discovery. Recall can assess the model’s ability to capture NLSs, while Hit-aPC further evaluates the percentage of NLSs captured. Additionally, the exploration cofactor is a variable that determines the rate of residues used to generate key fragments for each prediction.

The first parameter is whether to filter single residue. The principle of A2KA is to first extract important residues from the whole sequence, and then use the KSG (Key Segment Generation) algorithm to examine their distances with each other and generate segments. However, if a single residue can’t find another one close enough in sequence distance to form a segment, it will be left isolated, especially when the module uses a low exploration cofactor. Considering that an NLS segment generally has a minimum length of 3, we apply a random stretch on these single residues to form a segment; otherwise, these residues are ignored by our model. Another important parameter is whether to stretch the segments in a predefined way. When stretching, the model further extends the recommendation segment to a predefined length with a certain probability. Both strategies stem from our hypothesis that a residue or segment with high attention weight generated by A2KA not only indicates a high probability being an NLS but also suggests that the surrounding area is important.

Figure 3-b and Figure 3-c show the Hit-aPC and recall for all NLSs in the INSP datasets and NLSs of proteins in the Swiss-Prot database. The results of both datasets show the same tendency: stretching segments can bring improvement for Hit-aPC, especially for a low exploration cofactor. In the INSP datasets, the max improvement is about 0.1. For experimentally validated NLSs in Swiss-Prot, it is about 0.08. This improvement will be weakened as the exploration cofactor increases. In addition, stretching segments can also help achieve a better recall. While stretching segments benefits NLS localization, it can introduce more noise into recommendation sets, which consist of other important motifs such as DNA-binding.

On the other hand, whether to filter single residue has a significant impact on the recall. As shown in Figure 3-b, c, choosing not to filter single residue significantly improves recall and Hit-aPC. For both INSP datasets and experimentally validated NLSs in Swiss-Prot, not filtering single residue brings a maximum recall and Hit-aPC improvement of 0.25 and 0.18, respectively. Furthermore, the model approaches a recall of 1.0 earlier. For the INSP dataset, NLSExplorer without filtering achieves a recall of 1.0 with a cofactor of 0.3, compared to 0.5 of NLSExplorer with filtering. For the experimentally validated NLSs in Swiss-Prot, NLSExplorer without filtering achieves a recall of 1.0 at a cofactor of 0.3, compared to 0.6 of NLSExplorer with filtering. This suggests that the residues extracted by A2KA are indeed some special points in NLSs. Stretching these isolated residues helps the NLSExplorer find NLSs overlapping with them. Furthermore, this implies that these extracted residues indicate not just the importance of the amino acids themselves, but also the area including these residues or close to them. This also explains why stretching the generated segment can improve both recall and Hit-aPC.

In addition to experimentally validated NLSs, there are various support levels of NLSs in Swiss-Prot. We collect these NLSs along with their support levels and test whether A2KA is still effective at capturing them (see the statistic and different support levels at Extended Data Fig. 3). We set the parameters of A2KA to not filter single residue and not stretch segments. Figure 3-d shows the detected results for NLS data with different support levels, they share the same overall tendency but exhibit local differences. The NLS data labeled with “ECO:0000256” have a higher recall than others with the same exploration cofactor, while NLS data with “ECO: 00000303” label has the worst recall. All support levels can reach a recall of 1.0 when the exploration cofactor is high enough but not over 0.5.

To further investigate the characteristics of A2KA, we apply A2KA to proteins with five different types of domains. Figure 3-e shows the recall of A2KA for five types of motifs on proteins. Except for NLS and NES, A2KA cannot achieve a high recall for the other three characteristic regions when the exploration cofactor is not close to 1.0 (i.e., recommending the entire sequence). When the cofactor is 0.3, the recall for NES, DNA-binding, tRNA interaction, and RNA cap binding is 0.94, 0.60, 0.54, 0.42, respectively. For NLS, the recall is 0.96, indicating that nearly all NLS data is included in the prediction result of A2KA, whereas nearly half of the tRNA interaction and RNA cap binding is missed. This suggests that in the current exploration cofactor range (0.1-0.5), A2KA has a high probability of missing correct prediction except for NLS and NES.

Figure 3-f shows the reciprocal value of the exploration cofactor when the recall for the five motifs reaches different levels, it ranks in the order of NLS, NES, DNA-binding, tRNA interaction, and RNA cap binding, particularly when the exploration cofactor is low. This rank pattern highlights that high-ranked areas such as NLS and NES have significantly higher values than others. It partially reflects the recommendation preference of A2KA in NLSExplorer. We attribute this to A2KA’s attention mechanism that is trained to find key area highly correlated with the subcellular localization of the nucleus. A protein containing NLS has a high probability of entering the nucleus. When training the model to predict nucleus localization, the key area of NLS is helpful for the model to ensure proteins located in nucleus. Therefore, it is reasonable to find key area like NLS in a high priority. The results in Figure 3-e and Figure 3-f validate this point. Apart from NLS, NES is also an important domain highly related to nucleus. It consists of specific amino acids that aid in protein export from the nucleus. Therefore, proteins with NES are also initially located in nucleus and subsequently require NES for export.

In addition to NLS and NES, the recognition of other specific motifs or domains can also assist the model in predicting the localizations of proteins. Although they may not provide strong evidence to directly determine nuclear localization, they still are beneficial for the model to predict protein localizations. Therefore, we test the recognition performance for three additional motifs: DNA-binding, tRNA interaction, and RNA cap binding.

DNA-binding mainly occurs in the nucleus and has been reported to overlap with NLS^29, 30^. However, there also exists DNA at mitochondria, such as mtDNA, which means DNA-binding can also happen in mitochondria. In eukaryote, tRNA is synthesized in the nucleus through the transcription of tRNA gene and is processed to form mature tRNA. tRNA interaction occurs in the nucleus, but the major function of tRNA is to transfer amino acids to the ribosome for protein formation. RNA cap binging occurs in the nucleus, cytoplasm or ribosome. The cap-binding complex formed in the nucleus can influence the function of mRNA to fine-tune protein expression^31^. Overall, these three types of motifs in proteins can reflect the protein localization, but they cannot definitely confirm that a protein is located in the nucleus. Therefore, A2KA will not prioritize these motifs highly, as shown in the model’s recommendation order in Figure 3-f. This also explains the consistent components of the segment recommended by A2KA and helps interpret what might be the noise when predicting NLS.

In addition, we evaluate the computational efficiency of A2KA and the recommendation system in NLSExplorer. Note that our model uses embedding generated by ESM1b-650M in the initial stage; the embedding generation time is shown in Extended Data Fig. 2. Since ESM can only accept protein lengths less than 1022, we choose sequences with length ranging from 3 to 1022 in Swiss-Prot as the input to A2KA and segment recommendation system to test their computational time on a single RTX 3090 GPU. Figure 3-g shows the computational time, with the maximum computational time for A2KA and the recommendation system in NLSExplorer being no more than 0.29s and 0.18s, respectively. The results show that the primary time consumption is from the ESM embedding generation, which takes around 11s.

### NLSExplorer highlights potential NLSs with high sequence and structure similarities

As demonstrated previously, A2KA is able to detect segments within a protein sequence that play important roles in the nuclear transport process, such as NLS. Additionally, we investigate how attention mechanisms help identify these key areas. We run NLSExplorer on a transcriptional regulatory protein called Autographa californica nucleopolyhedrovirus^32^ with Swiss-Prot ID P11138 and its MSA sequence with an E-value of zero from UniProtKB and Swiss-Prot. Figures 3-h and 3-i show the results of two groups for multi-attention display.

In the first group, P11138 with an HSP greater than 90% displays higher levels of attention around two patterns, “SA(A/T)GTKRK” and “(K/R)PK.”, NLSExplorer detects two segments containing these patterns with a high probability of being NLS. Additionally, the pattern “RPK” shows greater attention than “KPK” indicating a nuclear import tendency with a single amino acid variation. In the second group, the amino acid sequences ‘KRK’ and ‘RPK’ are also highly notable, as the segments containing these sequences are predicted to function as nuclear localization signals (NLS) in proteins A0A288Q7F6 and A0A172WZ84. In summary, P11138 and its MSA sequence with an E-value of zero are both predicted to have a high probability of containing NLS, suggesting that these nuclear virus proteins may rely on NLS to enter into the nucleus.

Along with the similarity in protein sequences, we investigate whether NLSExplorer could recommend potential NLS segments that share common structural characteristics. We run NLSExplorer on the nuclear proteins of Swiss-Prot and extract three amino acid segments with the highest cosine similarity based on their representations generated by NLSExplorer’s recommendation system. We use the 3D structures predicted by Alphafold^33^ as their structural references (see details in Supplementary File 2). We first analyze the overall structural similarity of the proteins containing potential NLSs. Figure 3-j shows the structural alignment, and Figure 3-k shows their sequence length and sequence similarity. The three proteins have sequence identities ranging from a maximum of 0.68 to a minimum of 0.44 with each other. Compared to the sequence identity, their structural similarity is higher, with TM-scores^34^ ranging from a minimum of 0.84 to 0.94 and RMSD (Root Mean Square Deviation) of 1.85 to 2.95. Furthermore, we extract the aligned NLSs. Among the three NLSs, although “AVTRTAAHTHIK” only shares 5 and 6 identical amino acids with “RIERIGAHSHIR” and “RTQRIAAHSHVK” out of a total of 12 amino acids, it has RMSDs of 0.99 and 0.89 with them (Figure 3-l), indicating high structural similarity. This aligns with the overall result that, while the sequence identity for these three proteins is not very significant, their structural similarity is remarkable. Moreover, the local similarity of the recommended NLSs shows a higher level of structural conservation compared to the overall sequence similarity, implying that this area may be a more conserved functional region within the protein.^35^

Figure 3-m showcases the structures of additional recommended NLSs with their near NLSs based on the cosine similarity in the recommendation system’s embedding space. The recommended NLSs exhibit high local structural conservation, even though the proteins containing them have different global structures. This result suggests that proteins with varying global 3D structures can rely on NLS fragments with highly conserved local structures for nuclear localization.

Figure 3-n illustrates some interesting cases where four recommended NLSs from two different proteins cluster closely together and show high structural similarity. In each protein, two structurally identical NLSs coexist. Further investigation reveals that these NLSs belong to DAZ4_HUMAN (Deleted in azoospermia protein) and DAZ1_HUMAN, both of which are RNA-binding proteins essential for spermatogenesis ^35^. Given the susceptibility of the DAZ gene to deletion, we speculate that this may represent a novel mechanism where organisms use repeated NLSs to enhance the efficiency of protein entry into the nucleus. This would potentially mitigate the negative impact on spermatogenesis caused by the loss of these proteins for the reproductive process due to the increasing occurrence of structurally isomorphic NLSs.

### Attention Module Analysis in A2KA for Optimizing NLS Detection with Selective Strategies

Language models provide biological insights by incorporating the knowledge captured during pretraining into the generated protein embeddings. The Basic Attention Units (BAUs) in A2KA responds to various input contexts by focusing on two key aspects of the protein embedding: how to effectively aggregate amino acid vectors into an overall representation, and which features of a vector play a more significant role in different tasks. To address these aspects, refining the protein embedding using attention mechanism can help aggregate information to closely align with the current task. BAUs introduce row attention and column attention (Figure 4-a) to extract useful information and filter out irrelevant elements from the entire protein embedding (Methods).

**Figure 4.**
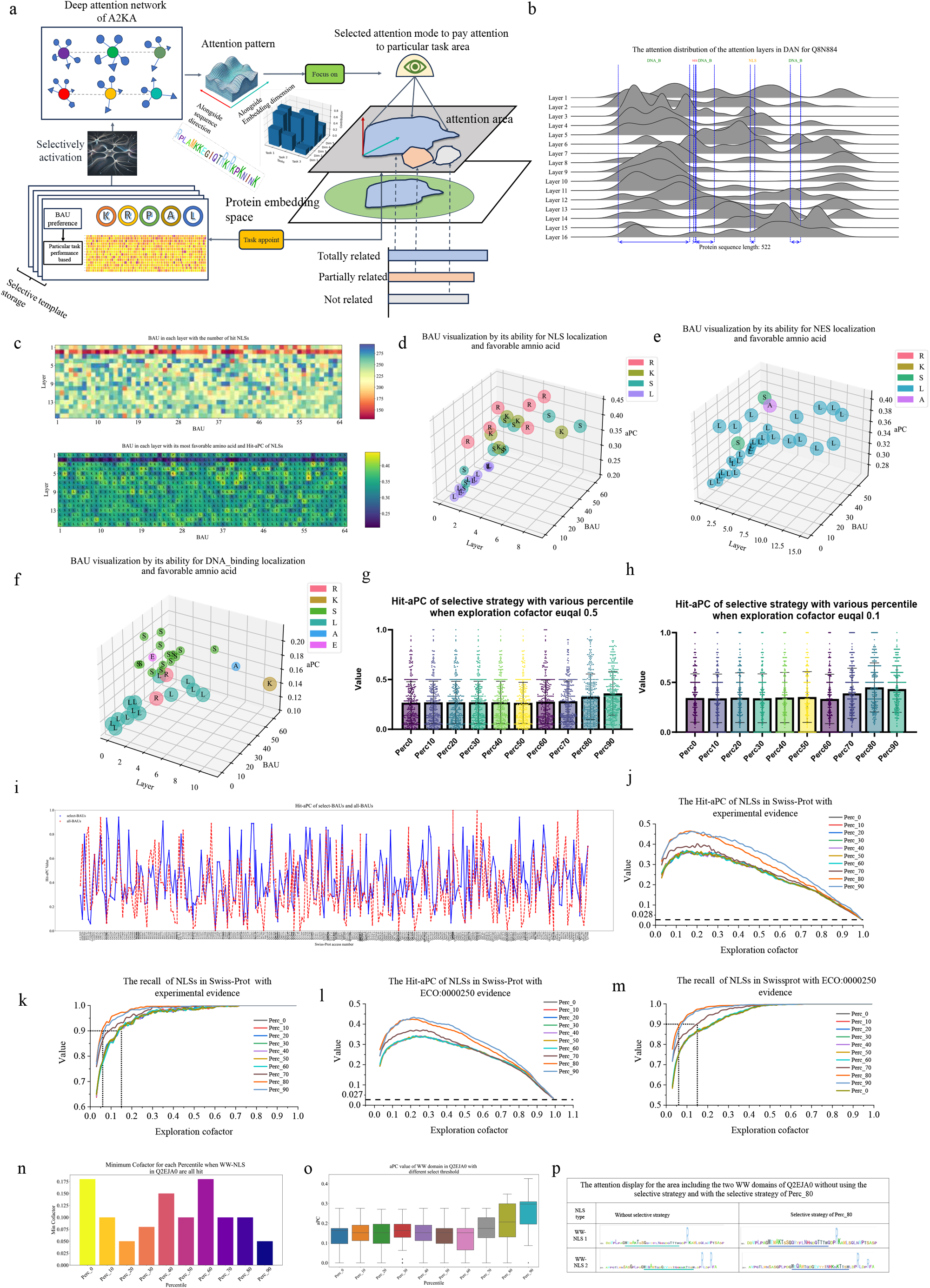
In-depth exploration of the attention mechanism for A2KA. **(a)** The BAUs (Basic Attention Units) in A2KA utilize row attention and column attention to focus on both the protein sequence and embedding dimensions, effectively extracting key information that is related to current task. The introduction of a selective strategy further enhances the A2KA module by activating well-performing BAUs and deactivating poorly performing ones. **(b)** The attention distribution of the attention layers in DAN (Deep Attention Networks) for the proteins with Swiss-Prot ID Q8N884. **(c)** The Hit-aPC and hit numbers of BAU for NLS in Swiss-Prot with experimental evidence. **(d)** Top 20 and last 20 hit numbers of BAU for experimentally validated NLSs in Swiss-Prot, the size of points means the hit number of BAU. **(e)** Top 20 and last 20 hit numbers of BAU for experimentally validated NESs in Swiss-Prot. **(f)** Top 20 and last 20 hit numbers of BAU for experimentally validated DNA binding site in Swiss-Prot. **(g)** The performance comparison using different selective strategies for experimentally validated NLSs in Swiss-Prot with an exploration cofactor of 0.5. **(h)** The performance comparison using different selective strategies for experimentally validated NLSs from Swiss-Prot with an exploration cofactor of 0.1. **(i)** The recommendation result for 298 NLSs from Swiss-Prot with experimental evidence when using “Perc_80” and all BAUs, Hit-aPC of 0 means this NLS is not a hit. “Perc_80” means selecting BAU with the performance greater than 80% of the optimal performance.**(j)** The Hit-aPC of various thresholds for the selective strategy on NLS data with experimental evidence. **(k)** The recall of various thresholds for the selective strategy on NLS data with experimental evidence. **(l)** The Hit-aPC of various thresholds of the selective strategy on NLS data with “ECO:0000250” evidence. **(m)** The recall of various thresholds of the selective strategy on NLS data with “ECO:0000250” evidence. **(n)** The minimum exploration cofactor when A2KA with different selective thresholds can locate the two WW-NLS for protein YAP1 (Swiss-Prot ID Q2EJA0). **(o)** The Hit-aPC distribution for protein YAP1 when utilizing different selective thresholds as the exploration cofactor changes from 0 to 1. **(p)** The attention distribution comparison of A2KA without the selective strategy and A2KA with the selective strategy of “Perc_80”. The left subfigure uses attentions from all BAUs, and the right subfigure uses “Perc_80” to aggregate attentions.

During model training of NLSExplorer, the attention mechanism within A2KA is shaped into a specific module tailored to nuclear localization prediction tasks. As a result, various BAUs from different layers exhibit distinct attention convergence patterns in different regions (Extended Data Fig. 4). Some BAUs excel at NLS discovery, while others perform poorly in this task but show strengths in others, such as DNA-binding site detection. These traits are reflected in their preference to specific amino acids. In this section, we explore the characteristics of BAUs in depth and introduce a selective strategy to further enhance the prediction performance in specific tasks.

### In-depth investigation of BAUs in A2KA

We initiate our inquiry on a particular class of proteins, CGAs (Cyclic GMP-AMP synthase), whose Swiss-Prot ID is Q8N884. This protein contains DNA-binding region, NLS, and NES. These multidimensional characteristics make it a very special protein. It interacts with poly-ADP-ribosylated (PARP1) to prevent the formation of PARP1-Timeless complex in the nucleus, which has a suppress effect on homologous recombination of DNA and thus promote the progression of the tumor^36^. It also plays an important role in innate immunity by functioning as a key DNA sensor when binding to double-stranded DNA in cytoplasm to trigger type-I interferon production^37^. We examine the attention distribution for BAUs in A2KA and extract the attention weights generated by each BAU for each amino acid in proteins. Then, the attention from individual BAUs is grouped by the layer index of DAN and summed together for visualization. Figure 4-b shows the visualization of layer attention distribution for CGAS. Interestingly, for the DNA-binding region in the C-terminal of the protein, which has the longest length, there are a large number of 21 Wave crests, indicating that A2KA places a lot of attention on it. The other two DNA-binding regions are also included or close to the four attention wave crests, suggesting that the model focuses on these areas when making predictions. For the NLS region, although it is far smaller than others, there are still eight wave crests in or close to it, which shows a high attention priority for NLS. As for NES, despite its much smaller length compared to other specific regions, the model shows two peaks of attention inside it. The layer attention visualization for CGAs illustrates that each attention layer of DAN has diverse focus patterns, and for special area in proteins, these diverse focus patterns can assume different roles to complement each other to comprehensively capture different types of important regions.

In the previous layer attention visualization, we observe that not every layer contributes to capturing important areas like NLS. On the contrary, some layers have not yet focused their attention on the NLS area. From an alternative perspective, it is natural to suppose that not all attention units in A2KA are responsible for NLS detection.

We further investigate the attention mechanism of A2KA in a finer granularity. We extract all of the 298 NLSs with experimentally validation evidence from Swiss-Prot. We employ a parameter set without filtering single residue and without stretching segment using an exploration cofactor of 0.3. We extract the attention distribution of each BAU(16 layers×64 units) and input them to the segment generation algorithm to obtain recommendation sets. Subsequently, we collect the most frequent amino acids in these recommended segments as the favorite amino acids for each BAU. Then, we count the total numbers of NLS hit by NLSExplorer and calculate Hit-aPC of NLS for each BAU.

Figure 4-c displays the favorite amino acids of each BAU and their total hit numbers. BAUs in A2KA show diverse amino acids preference such as K, R, L, and S. For the tasks of NLS detection, we select the top 20 and the bottom 20 BAU based on the rank of the hit number and show their favorite amino acids and Hit-aPC in Figure 4-d. The BAUs with favorite amino acid of K and R occupy a large proportion of the top 20 BAUs and are not present in the last 20 BAUs. It implies that BAUs pay attention to segments rich in amino acid K and R are more likely to achieve superior performance for NLS localization prediction task. This aligns well with the established knowledge that NLS generally consists of KR-enriched peptides^3^. Figures 4-e and 4-f display the top 20 and bottom 20 amino acid preferences for the tasks of NES (Nuclear Export Signal) and DNA binding site detection. Compared to NLS prediction tasks, the BAUs that perform better on these two tasks exhibit significant differences in their amino acid preferences. This result indicates that the attention patterns shaped by the nuclear-related tasks in BAUs are various, making them adaptable to different, more specialized tasks.

### The selection strategy based on BAUs enhances A2KA’s ability to explore potential NLSs

Based on the results that the BAU with different favorite amino acid has a huge difference on the NLS prediction performance, we suppose that for a specific task such as NLS prediction, it is advantageous to selectively adopt BAUs with particular preference rather than using all BAUs. To validate this hypothesis, we implement a selective strategy to pick BAUs based on their performance on NLS data with experimental evidence. Then, the attention distributions of selected BAUs are aggregated for key segment generation. We divide the intervals and names of the BAU group based on whether the BAU achieves the assigned percentage of optimal Hit-aPC and hit numbers. For example, “Perc_80” means the BAUs in this group all have Hit-aPC and hit numbers exceeding the 80% of the optimal Hit-aPC and hit numbers. We evaluate the impact of the selective intensity on Hit-APC by varying the percentile from 0% (no selective strategy) to 90%. Figures 4-g and 4-h compare the results under the exploration factors of 0.1 and 0.5. Both Perc_80 and Perc_90 show significantly better overall Hit-APC than the other percentiles, where Perc_80 performs better at lower exploration factors and Perc_90 excels at higher ones. This suggests that selecting BAUs for specific tasks can enhance the overall model’s performance. Figure 4-i shows the recommendation results of A2KA for 298 NLSs with experimental evidence using “Perc_80” strategy with exploration cofactor equal to 0.3 compared to selecting all BAUs. In some cases, such as “Q76U48” and “Q13177”, selecting all BAU fails to capture the correct NLSs for these two proteins. In contrast, A2KA with the selective strategy “Perc_80” achieves Hit-aPC values of 0.5 and 0.625 for NLSs of the two proteins, respectively, compared to selecting all BAUs (Hit-aPC of 0). The introduction of this selective strategy improves the overall Hit-aPC and successfully identifies 10 previously uncaptured NLSs. This demonstrates that the introduction of BAUs that are not good at detecting NLSs may has a negative impact on the prediction performance.

We further conduct experiments on NLS data to investigate the impact of the selective strategy’s threshold on the exploration results of A2KA. Since the selection is according to the prediction result on NLS data with experimental evidence; we also apply this approach to NLS data with ‘ECO:0000250’ evidence to obtain more robust results. In Swiss-Prot, we have collected 2005 entries. Figure 4-j, k, l, m shows the Hit-aPC and recall on NLS with the experimental evidence and “ECO:0000250” evidence. For both datasets, introducing the selective strategy improves the A2KA, significantly boosting the recall and Hit-aPC with the maximum improvement of over 0.1. In addition, the model’s performance improves notably when the selection percentage is higher than 70%, and the improvement is marginal when it is lower than 70%. From the perspective of an overall performance, A2KA with Perc_80 and Perc_90 strategies both achieve the best results. A2KA with Perc_90 yields a slightly higher Hit-aPC, whereas a lower recall than A2KA with Perc_80. In addition, the recall curve shows a consistent upward trend with increasing exploration cofactor, the Hit-aPC increases until it reaches a maximum value with an exploration cofactor of 0.2, then declines. We can see that as the exploration cofactor increases, it brings more possibility for the model to find NLSs. However, it also brings more possibility for introducing other areas that may be noise for the final NLS localization prediction.

We evaluate A2KA of NLSExplorer on a novel type of NLS recently discovered by Yang^38^, characterized by a unique globular structure known as WW-NLS. This WW-NLS variant, identified within folded WW domains, has been experimentally confirmed to serve as an NLS in YAP1. For our investigation, we utilize the YAP1 protein sequence containing the validated WW-NLS motifs to assess the capability of NLSExplorer in detecting and locating these emerging NLS variants. Figure 4-n shows the minimum exploration cofactor required for the two types of NLS detection, Perc_90 and Perc_20 has the best performance with an exploration cofactor of 0.04 to capture all the NLSs. In contrast, A2KA with Perc_0 (which means selecting all the BAUs) and Perc_60 need the exploration cofactor of at least 0.175 to detect all NLSs. Figure 4-o shows the Hit-aPC score of these two NLSs when the exploration cofactor varies from 0 to 1. It shows the same tendency with our large-scale experiment on NLS data from Swiss-Prot. When using a strategy with a percent greater than 70, the A2KA model achieves better performance. In addition, we visualize the attention change when applying the ‘Perc_80’ strategy in Figure 4-p. It shows that the introduction of the selective strategy directs the attentions to focus on the WW domain, particularly highlighting the amino acid tryptophan (‘W’) at the start of this area. Furthermore, there is also high attention (amino acid “R”) around these two domains, which implies a high probability of the surrounding area, including WW-domain, for being NLSs. The result shows the exploration ability of NLSExplorer and demonstrates its ability in recognizing novel NLS regions in the protein sequence.

### NLSExplorer for NLS Discovery and Pattern analysis in Swiss-Prot

Understanding the fundamental principles of amino acid formations is crucial for developing NLS-based nuclear complexes for medical applications and mechanistic research on nuclear transport. Beyond making single predictions for individual proteins, we leverage NLSExplorer’s effectiveness in accurately identifying NLSs and its strong discovery capabilities for nuclear transport-related segments to comprehensively conduct the investigation into their presence, pattern characteristics and cross-species properties.

We use NLSExplorer to extensively explore the NLS universe across all experimentally validated nuclear proteins in Swiss-Prot. The resulting key segments are used to construct a map for exploring the NLS space across four distinct levels (Figure 5-a). The discontinuous patterns of NLSs, characterized by the dynamics and stability of amino acids, help identify the amino acids that play the most critical roles in NLS segments. The nuclear transport map employs hydrophobicity and entropy as measurement and projection baselines, with the recommended fragments orderly to highlight the unique features of transport segments for different species. A candidate NLS map is also created to visually display the overall distribution of recommended NLS segments, equipped with a convenient search function that serves as a valuable reference for NLS research. Lastly, a species correlation map is developed using TF-IDF(Methods) as quantification approach for the segments. Its graphical visualization reveals the connections between nuclear transport segments across species, partially uncovering evolutionary correlations among them.

**Figure 5.**
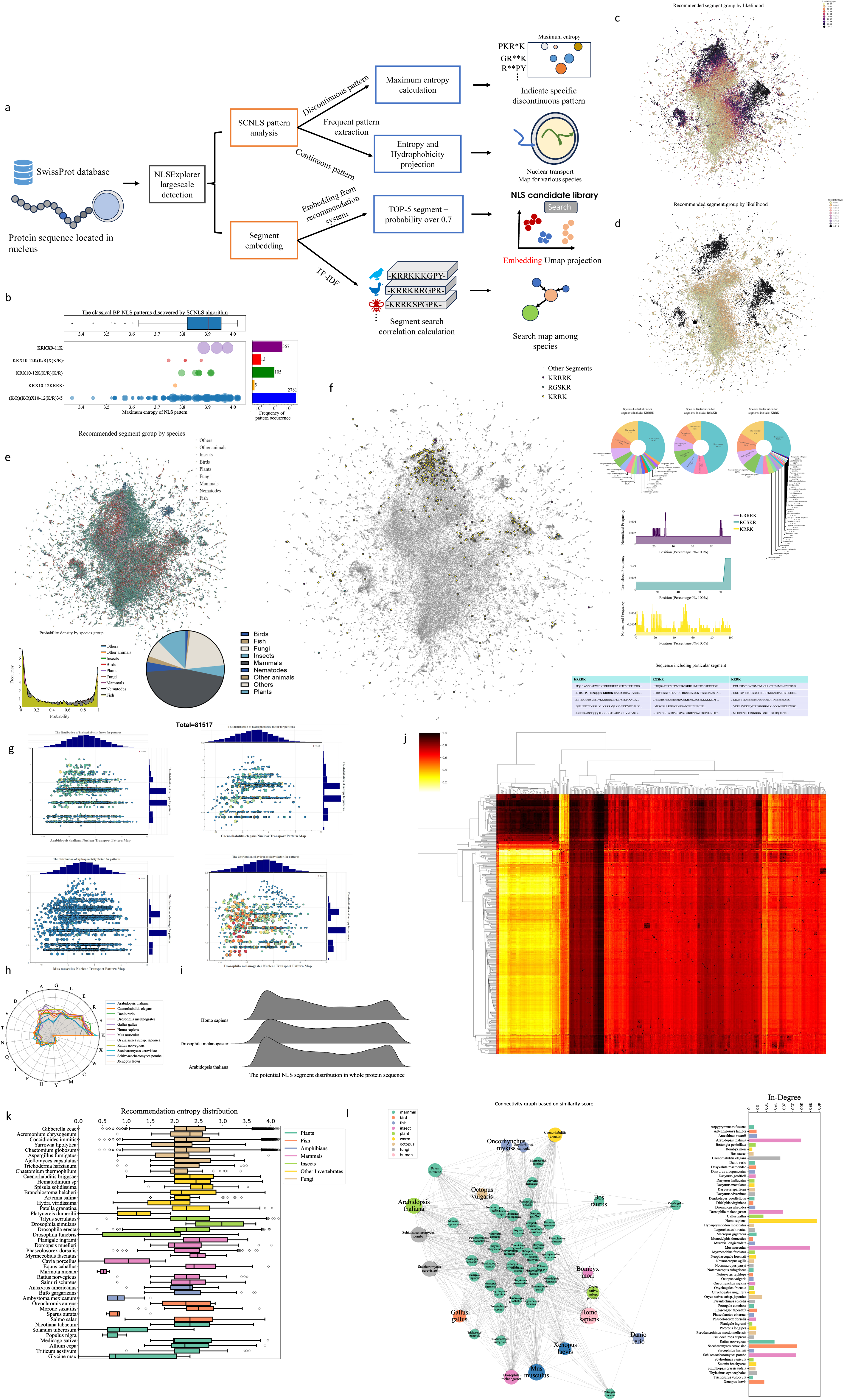
The NLS pattern and candidates revealed by NLSExplorer-SCNLS algorithm. (**a**) We use NLSExplorer to explore the NLS universe and construct the maps from four different perspectives. **(b)** The Bipartite patterns of NLS discovered in SeqNLS is detected by SCNLS, X10-12 means 10 to 12 amino acids that can vary to any type, (K/R)3/5 represents there are at least 3 amino acids of being K or R in five amino acids. The x-axis represents the maximum entropy of the NLS pattern, which reflects the diversity and uncertainty of NLS patterns. The size of bubbles indicates the occurrence of different patterns. **(c)** The recommended segments of experimentally validated nuclear proteins annotated by their probability of being NLSs. **(d)** The recommended segments of experimentally validated nuclear proteins are annotated by their probability of being NLSs, within the ranges of 0.0 to 0.2 and greater than 0.9. **(e)** The recommended segments of experimentally validated nuclear proteins annotated by their species. **(f)** The recommended segments of experimentally validated nuclear proteins annotated by the searching segment contained in them. **(g)** The Nuclear Transport Pattern Map for various species. x-axis represents the hydrophobicity for a specific pattern and y-axis indicates the max entropy. We set the point sizes of a pattern based on its occurrence. **(h)** The amino acid occurece in various species, we select the species with more than 100 protein sequences located in nucleus. **(i)** The potential NLS segment distribution in the protein sequences in Homo sapiens, Arabidopsis thaliana, Drosophila melagogaster, they are representive species of mammals, insects, plants in the NLS research field. **(j)** The searching correlation hierarchically clusters the heatmap among 416 species. The rows of this map indicate the query species, while the columns represent the species to be searched against. **(k)** The entropy distribution for the recommendation segments amnog the species, we randomly chose 50 species to show and group them by categories. Categories with fewer than three species are filtered out. **(l)** The species search map built on the search correlation score. We choose in-degree more than 20 to visualize. The in-degrees of species are represented as the size of points.

### NLSExplorer-SCNLS facilitates the investigation of NLS patterns

The continuous pattern of NLSs can be easily identified by tracking the occurrence of substrings. However, this method is not able to identify the frequent pattern with gaps. We propose SCNLS algorithm to explore the discontinuous pattern of NLSs. We use NLSExplorer to mine potential NLS in the Swiss-Prot database. We set the parameters of A2KA in NLSExplorer without filtering single residue and stretching segments with an exploration cofactor of 0.3. Next, we extract 14283 protein sequences from Swiss-Prot with experimental evidence for nucleus localization. To address the length limit of ESM embedding, we cut the protein sequence of over 1022 into several parts, ensuring each part is less than 1022 amino acids. When we recommend potential NLSs, these sub-proteins are treated as independent entities. We then apply the selective strategy of “Perc_80” for A2KA in NLSExplorer to recommend potential NLSs for each protein and sub-protein. Subsequently, we use SCNLS to analyze the set of potential NLSs to identify discontinuous NLS patterns.

In order to confirm the effectiveness of the discontinuous NLS pattern mined by NLSExplorer-SCNLS, we validate our results using classic discontinuous Bipartite (BP) patterns of NLS. We extract all four BP patterns in the SeqNLS^39^, which is an excellent predictor with a relative high precision on the test set of INSP^27^.Figure 5-b shows the results of SCNLS and BP patterns of SeqNLS. Numerous BP patterns are enriched with (K/R)(K/R)X10-12(K/R)3/5, which is the most frequently occurring pattern with a frequency of 2781. Moreover, (K/R)(K/R)X10-12(K/R)3/5 is also the pattern template used by SeqNLS. The other three patterns are all detected by SCNLS, with the occurrence of 5 for KRX10-12KRRK, 105 for KRX10-12K(K/R)(K/R), and 13 for KRX10-12K(K/R)X(K/R). We measure the distribution range of the maximum entropy for these four BP patterns, and discover that they are generally within the range of 3.8-4.0. Furthermore, we extract the NLS patterns of the maximum entropy with the interval of 3.8-4.0. The most frequent pattern, KRX9-11K, is shown in Figure 5-b. KRX9-11K has three significant patterns and a total frequency of 357. In addition, it is a type of K/R rich pattern, consistent with the previous NLS patterns with a high probability of being potential BP patterns.

### NLSExplorer explores NLSs and nuclear transport segments in Swiss-Prot database

To advance the investigation and identification of NLS, we put all the potential NLSs recommended by NLSExplorer from the experimentally validated nuclear-proteins in Swiss-Prot into an interactive visualization map - NLS Candidate Library (Extended Data Fig. 5). Each point represents a recommended segment with the possibility predicted to be an NLS. Their coordinates are obtained by projecting the embedding from the recommendation system of NLSExplorer using UMAP. In this map, NLS is labeled with the probability predicted by NLSExplorer, selecting a threshold can filter segments to visualize NLS in various confidence levels. Figure 5-c shows the points labeled by their NLS likelihoods. The darker areas represent points with higher probabilities, while the lighter areas indicate those with lower probabilities. Figure 5-d further highlights the regions with probabilities greater than 0.9 and the range of 0.0 to 0.2. These demonstrate a clear stratification pattern, where high-probability points cluster into three main regions. The remaining points fill the gaps between these clusters as intermediaries with probability increasing as one moves closer to the groups. This suggests that, compared to the darker regions, predicted to have a high likelihood of being NLSs, the lighter areas may represent potential new types of NLSs that were not included in the training dataset of the recommendation system. These regions could serve as promising areas for future investigation. In addition, analyzing them from a species-based perspective helps reveal distributional differences in NLS probability density and suggests the scale for NLS data (Figure 5-e). This map also includes a search function. By inputting amino acids segment, all potential NLSs containing that segment are highlighted, and relevant information, such as the species containing them, their detected positions, and their location within the full protein sequence, can be retrieved (Figure 5-f).

We develop another interactive map called Nuclear Transport Pattern Map to visualize the potential continuous segment patterns mined by SCNLS algorithm. These segments for pattern analysis are obtained by running A2KA with the selective strategy ‘Perc_80’. As a result, the model maintains an optimal ability to detect potential NLS segments and other segments that may be important for protein nuclear transport, while keeping a low probability of introducing irrelevant segments. Figure 5-g shows the pattern map for various species, The differences in the properties of transport segments across species are characterized by variations in hydrophobicity scores and entropy distribution. Figure 5-h shows the amino acid occurrence of recommended segments for various species, revealing an overall enrichment of K, R and S. In Figure 5-i, we visualize the positions where they are located, the typical regions near the N-terminal and C-terminal of a protein correspond to the positions where an NLS is frequently observed.

Using the search correlation (Methods) as a standard to assess the segment relationship among species allows to explore the patterns and tendencies. Figure 5-j displays the search correlation cluster map for 416 species with experimentally validated nuclear proteins. There are several clusters for sepcies searching and being searched. For example, Octopus vulgaris is in a search cluster that consistently exhibits a high search correlation score with other species (See details in Supplementary File 2.). We further explore the characteristics of nuclear transport segments among species. The entropy distribution of recommendations among species shows a diversity. Some categories like fungi showing the overall coherence with the certain species contain several high-entropy segments, indicating a substantial diversity (Figure 5-k). In addition, we visualize this search correlation by building a search map in Figure 5-l. This map shows that the search tendency converges towards species with rich segment patterns like Homo sapiens. Interestingly, mammalian categories display a mutual search correlation among species and are centrally connected to other species frequently. In general, the segment characteristics among species exhibit interspecies difference and shared characteristics of potential nuclear-mediated segments in Swiss-Prot.

## Discussions

Although understanding of NLS is a long-term basic biological problem, developing NLS prediction models from limited experimental data poses big challenges. Models specifically dedicated to NLS prediction face challenges. Due to the limited types and insufficient quantity of NLSs used for training, they exhibit poor generalization. To address the challenge, we present NLSExplorer, which is a robust framework for NLS prediction and exploration. NLSExplorer is developed based on the biology knowledge enriched within protein embeddings generated from large language models, where the attention mechanism acts as the bridge to eliminate the gap among various prediction tasks.

The large language model can capture intrinsic correlations and biological knowledge with massive pre-training sequences through self-supervised learning^40^, It presents big opportunities for exploring and identifying key functional regions, specific structural features, mutation sites, and other crucial areas on protein sequences. NLSExplorer utilizes A2KA to establish connections between protein nuclear localizations and language models, effectively transferring pre-trained representations and nuclear localization information to the task of predicting NLSs through attention mechanisms. By leveraging the knowledge from pre-trained representations, the NLSExplorer opens the door for exploring NLS space. NLSExplorer is able to discover new types of NLS and reduce its dependence on the size of the training dataset, resulting in superior generalization performance.

Unveiling the properties of the predicted fragments offers valuable insights for advancing NLS research. The NLS candidate library is built by leveraging the exploration capabilities of NLSExplorer and includes not only known NLSs but also potential ones. The similarity between sequences and structures is reflected in the neighborhood’s relationship of this map based on the cosine similarity of segment embeddings. This helps build a comprehensive landscape for each prediction by automatically searching for the nearest neighbors and provides customizable parameters to meet various usage requirements. NLSExplorer-SCNLS provides a powerful tool to highlight the core amino acids of NLSs and discover discontinuous NLS patterns. It facilitates NLS template finding and uncovering novel types of NLSs patterns. The Nuclear Transport pattern map, mined by the SCNLS algorithm, provides a reference for potential NLS patterns and other key segments important for nuclear transport. The map helps analyze the potential NLS characteristic and uncover the evolution relationship of NLSs among species. In addition, it offers the possibility of promoting advancements in various applications like targeted drug delivery, novel treatments for nuclear-protein-related diseases, and the development of new nuclear proteins for biological research.

NLSExplorer can be further enhanced in several directions. The “basic attention unit” in A2KA only considers the information derived from the sequences, neglecting the structural information. Notably, recent advancements in structural embeddings^41-45^ have been observed. Incorporating structural information into the A2KA framework could aid in gaining inspiration from structural information, potentially improving the prediction performance. Another limitation lies in the computational complexity of the SCNLS algorithm, which suffers to high complexity in mining the NLS segments. Consequently, substantial computational resources are required to fully exploit the algorithm’s capabilities. Further refinement of the SCNLS algorithm to achieve higher computational efficiency can enhance the efficiency for pattern mining. In addition, the A2KA framework provides a general framework that help explore the potential domains or motifs important for subcellular localization, our next step involves in continuous pushing our exploration on more important areas.

## Methods

### Datasets for NLSExplorer Training and Validation

NLSExplorer consists of two major modules. The first module, A2KA (Attention to Key Area), leverages attention mechanism to generated candidate NLSs. The second module is the NLS recommendation system, which recommends NLSs from the candidate set generated by A2KA. In the initial training phase, the goal is to use the nucleus localization labels as a supervised signal to enable the attention mechanism to identify the amino acid segments associated with nuclear localization in the sequence. The statistic for all datasets is shown in Extended Data Fig. 3.

#### Training dataset for A2KA and NLS exploration

We collect all protein sequences located within the nucleus with experimental evidence. We collect 14283 proteins of 416 species from Swiss-Prot (2023_05) as positive samples. To build a balanced training dataset, we keep the numbers of positive samples and negative samples equal. Thus, we select an equal number of 14283 protein sequences from Swiss-Prot that are not localized within the nucleus (528,724 proteins) as negative samples. We denote this dataset NLSExplorer-p. Moreover, the nuclear proteins in NLSExplorer are used by our model to detect potential NLSs and construct the NLS Candidate Library.

#### Validation datasets

To evaluate the performance of NLSExplorer for predicting NLS, we use the INSP dataset^27^ as a rigorous benchmark set. INSP dataset is built from the latest nlsdb^46^ database, SeqNLS^39^ and Swiss-Prot. It includes a training set and hybrid test sets, each enriched with proteins from a diverse range of species. In addition, a specialized yeast test set is constructed. The training set, consisting of proteins from various species, serves as foundational data for model training to enable the model to predict NLS from different species. Yeast is a species frequently used for studying NLS characteristics, and it has the richest experimental validated NLS data across species. The yeast test set is designed to assess model’s ability to predict NLS with NLS data from particular species. On the other hand, the hybrid test sets, characterized by diverse species, enable a thorough examination of the model’s generalization across different species.

#### Various support levels for NLSs

In addition to NLS data supported by ‘ECO:0000269,’ we extract NLS data supported by other levels of evidences from Swiss-Prot 2024_3. In Swiss-Prot, ‘ECO:0000255’ indicates information generated by sequence analysis programs and confirmed by a curator. ‘ECO:0000303’ refers to information retrieved from scientific articles without experimental support. ‘ECO:0000305’ involves manually curated information by curators based on their scientific knowledge. ‘ECO:0000256’ represents automated information generated by the automatic annotation system of UniProtKB.

#### Characteristic domains recognition datasets

To test the ability of NLSExplorer to detect characteristic areas, we extract proteins with labels for DNA-binding, tRNA interaction, and RNA cap binding, from Swiss-Prot 2024_3 with ‘ECO:0000269’ evidence, indicating they are experimentally validated. The detail for constructing this dataset is provided in Supplementary File.

### Evaluation Metrics

We use recall, precision, *F1* score and *aPC* as evaluation metrics of the prediction model. *recall* indicates the model’s ability to fully catch the actual NLS data, while precision examines the correct rate of the model prediction. The *F1* score evaluates the overall performance. In addition, *aPC* measures the extent to which the model captures for single NLS, it is calculated when a prediction segment overlaps with NLS, *Prediction* − *aPC* is a rigorous metric that tests the performance by comparing all the recommended segments with actual NLS for test datasets, it is used for the prediction task. *Hit* − *aPC* calculates the maximum aPC of segments from the recommendation set with the actual NLS and averages them to get the final result. It is used to assess the catchability for NLS in exploration tasks.

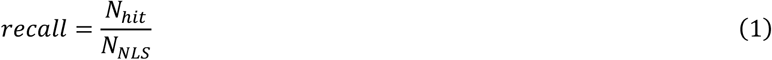

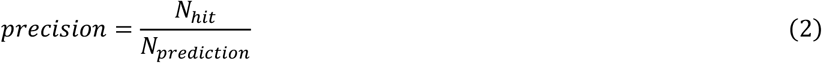

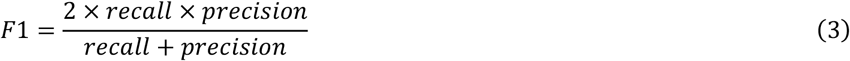

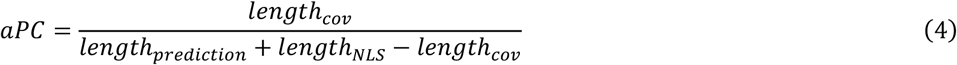

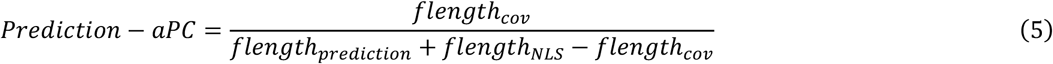

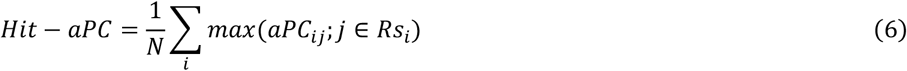

where *N*_*NLS*_ represents the total number of NLSs, *N*_*hit*_ indicates the number of NLSs detected by predictor, *N*_*prediction*_ refers to the total number of segments recommended to be NLS, *length*_*cov*_ indicates the overlapped length between a prediction segment and the actual NLS, *length*_*prediction*_ is the length of the prediction segment, *length*_*NLS*_ is the length of NLS, *flength*_*cov*_ means the total overlapped length of segments between the recommendation sets and the actual NLSs, *flength*_*prediction*_ is the total length of segments in recommendation sets, *flength*_*NLS*_ is total the length of NLSs, *i* is the index of NLS, *j* is the index of recommendation segments, *Rs*_*i*_ represents the recommendation segments set for NLS *i*.

### The Basic Structure of NLSExplorer

#### Attention to Key Area for key segment generation

##### Protein sequence task-specific attention

We introduce the concept of a protein sequence task-specific attention matrix (*A*) for a given prediction task. The representation of the sequence is *E*(*l*×*e*), where *e* represents the dimension of the embedding, and *l* is the length of the sequence. Note that *E* can be obtained by various methods; in this study, *E* is generated through the protein language model ESM. To determine the importance of amino acids at various positions, our model stores a query vector *q*(*e*×1). This vector forms an inner product with the vectors of amino acids in the entire sequence. A ReLU function is then applied to filter out negative values. The resultant values are subsequently processed by a softmax function to ensure the sum to be 1. After that, we utilize a scale factor, the square root of the sequence length *l*, as a divisor to mitigate the influence of the sequence length. Finally, *E* is aggregated from a matrix to a vector *v*_*t*_ according to the task-specific attention matrix *A*(*l*× 1). This vector is then input into a classifier to obtain the final prediction result.

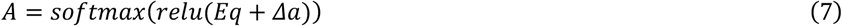

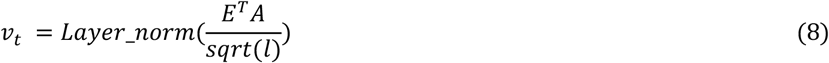

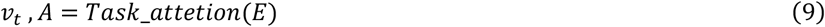

where the function *softmax* () means getting the softmax result along the length of sequence. *A* is the task-specific attention matrix, each value in *A* signifies the importance of residues in its corresponding position contributing to the prediction result. *q* is the stored query. *E* is the representation of the protein sequence. *l* represents the length of the protein sequence. *Δa* represents the bias vector. *sqrt*() denotes mean square root, and *Layer_norm* refers to the Layer Normalization. *relu* denotes ReLU function. *v*_*t*_ represents the classification vector generated by considering the attention weight of amino acids in different positions.

The introduction of ReLU function is to strengthen the representation of the residue with high (positive) attention values and weaken those with low (negative) attention values, making the attention mechanism more focused. NLSExplorer uses a large number of BAUs equipped with task-specific attention to form an attention network, which requires various units to be capable of capturing different sequence patterns. LayerNorm can normalize the generated vector *v*_*t*_ to maintain a stable distribution, thereby harmonizing the allocation of attention units. This balances the contribution of different attention units for the prediction result and prevents the final result from being dominated by a few specific BAUs, which makes the network robust and interpretable.

##### Attention-based Classifier Network Att_mlp

We employ a Multilayer Perceptron (MLP)^47^ as a linear classifier to handle various downstream tasks. For a protein representation *E*, we first use protein sequence task-specific attention to aggregate the amino acid representation in different positions to generate the vector *V*. This vector is then fed into an MLP to get the final result.

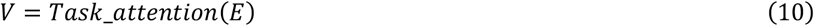

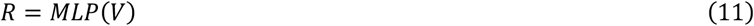

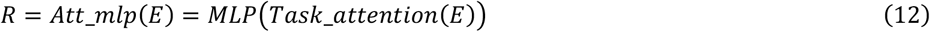

where *Task_attention* denotes protein sequence task-specific attention, *MLP* refers to Multilayer Perceptron, *R* is the prediction result, *Att_mlp* is the module that incorporates *Task_attention* and *MLP* to get the final result *R*.

#### Deep Attention Networks

We first devised a basic unit to form our exploration network. BAU employs a simple yet clear mechanism that focuses on using attention distribution to enhance and provide interpretation for specific tasks. As shown in Figure 6-a, BAU serves as the core component of the entire network. It is designed to enhance sequence representations for prediction and assist in model interpretation.

**Figure 6.**
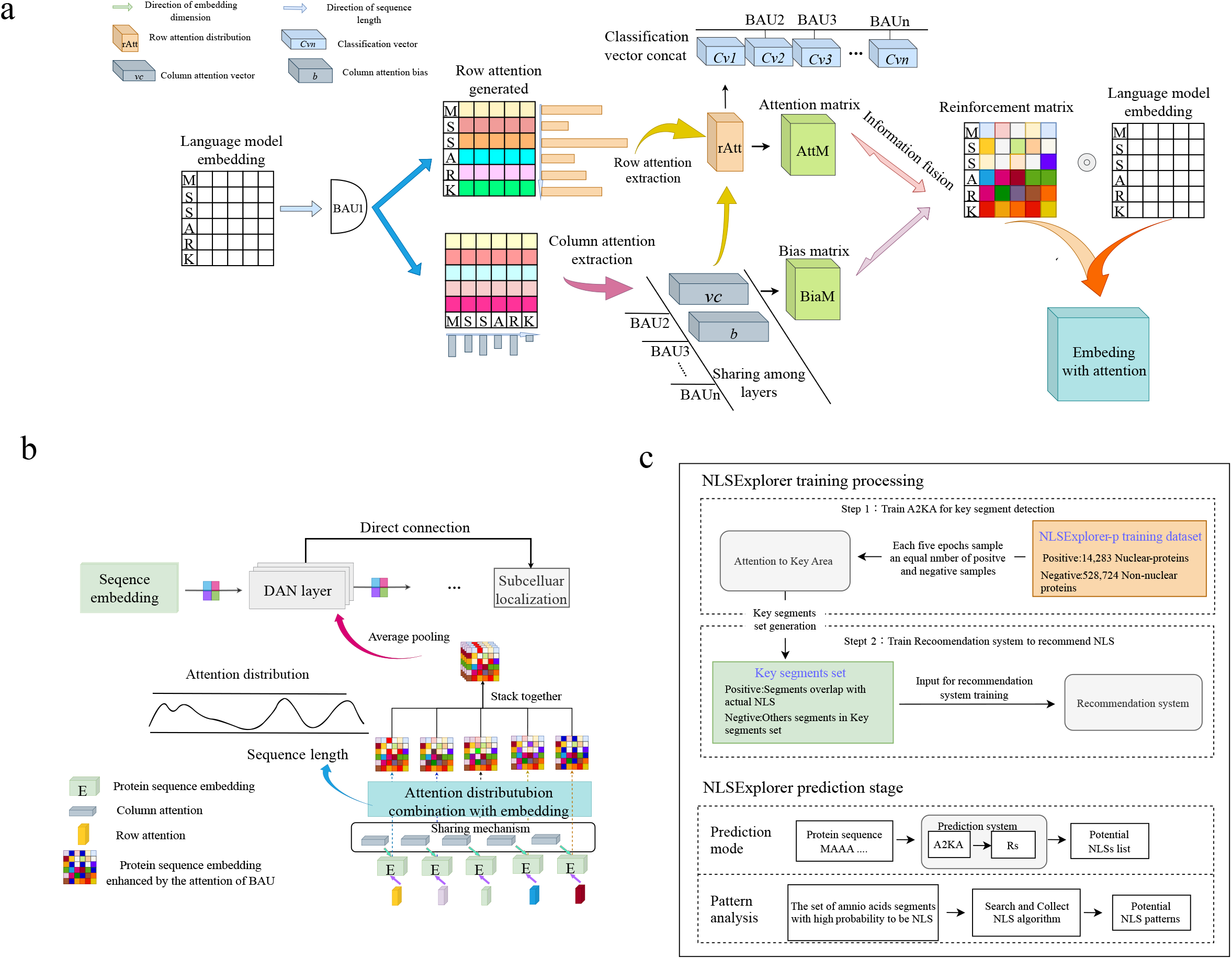
The basic structure and training details of NLSExplorer. **(a)** The “basis attention unit” is designed with row attention and column attention to generate row attention distribution maps and column attention distribution maps. The attention maps enhance the embedding, and the row attention map serves as a reference for key areas in the sequence. **(b)** The deep attention network is composed of attention layers; each is built upon the “basic attention unit.” **(c)** The introduction of training details and prediction stage for NLSExplorer.

##### Basic Attention Unit (BAU)

For a protein representation *E*(*l*×*e*), the BAU provides two kinds of attention mechanism to reinforce and highlight the important area. They are row attention and column attention, although similar to MSA transformer^48^, which have entirely different structures and functions.

###### Row Attention

Each row in a protein representation *E*(*l*×*e*) represents the vector representation of amino acids. As the name implies, row attention operates from the perspective of amino acids. It aims to utilize the attention mechanism to identify the amino acid important for specific tasks, thereby assisting the entire network in making predictions. In BAU, hence, we treat the task specific attention as the row attention:

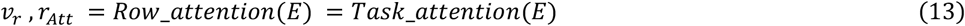

where *r*_*Att*_ (*l*×1) stands for the attention distribution map along the sequence, *E* is the representation matrix of the sequence, and *v*_*r*_ (*e*× 1) is the classification vector, which is the same as the formula 9.

###### Column Attention

Row attention can handle the importance distributed across the entire sequence. We want our model to also consider the varying levels of importance of features in different dimensions, which contribute to the final prediction result. Hence, we introduce a vector *v*_*c*_ (*e*×1) as the column attention vector. Values in different dimension of *v*_*c*_ are used to rescale the values in different dimension of the representation. In a protein representation *E*(*l*×*e*), each column indicates different features of the representation, so we refer to the second mechanism, which is focused on rescaling these features, as the column attention.

Building upon the distribution *v*_*r*_ formed by the row attention mechanism, an attention-enhanced matrix *A*_*r*_ (*l*×*e*) is constructed as: 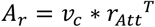. Additionally, we introduce a vector *b* to learn a current bias for each dimension of the embedding. The enhanced representation *E*_*r*_ is derived by performing the Hadamard product between the attention-enhanced matrix *A*_*r*_ and the overall sequence representation *E*.

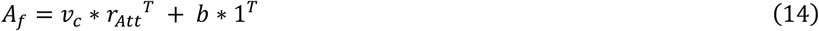

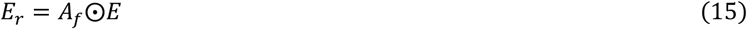

##### Deep attention layer based on BAUs

We combine *j* BAUs to form the attention layer (Figure 6-b). We generate row attention distribution from each BAU, and then the column attention unit is shared among BAUs.

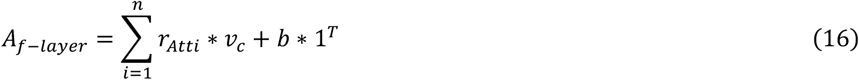

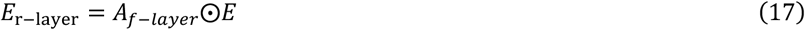

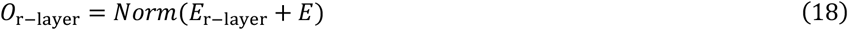

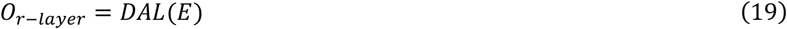

where *A*_*f*−*layer*_ is the an attention-enhanced matrix generated by deep attention layer, *r*_*Atti*_ is the row attention distribution of the *i-*th unit, *b* is the current bias for each dimension of the embedding, 1 indicates a vector with all dimension equal to 1, *v*_*c*_ is the shared column attention distribution, *E*_*r*−*layerr*_ is the representation enhanced by *A*_*f*−*layer*_, *E* is the embedding input, *O*_*r*−*layerr*_ is the output of the attention layer. *Norm* refers to the Normalization function, in this study we choose it as Layer Normalization. *DAL* denotes the deep attention layer.

##### Deep Attention Network

A Deep Attention Network (DAN, Figure 6-b.) is formed by connecting a defined number of deep attention layers. The attention mechanism in DAN needs to be formed by supervised tasks; therefore, we improve its structure to endow it with classification ability. In a DAN, the representation of the protein sequence *E* is first fed to this network and then hierarchically reinforced by the attention mechanism within the network. The final output of DAN, *E*_*f*_, is then fed to a protein sequence task-specific attention to get a classification vector *v*_*f*_. Additionally, the row attention in each BAU generates the classification vector *v*_*r*_, we use *v*_*r*_ from different BAUs in different layers to make prediction. The design is similar to the residue skip^49^; for a DAN with many layers, excepting the final classification vector *v*_*f*_, it provides another short and direct way for the model to train the mechanism in BAU by using *v*_*r*_ to influence the prediction result. This approach allows the gradient to backpropagate more directly, making it easier to adjust the attention module for each BAU. Considering all these aspects, the final prediction and the integration attention distribution map *r*_*t*_ can be obtained:

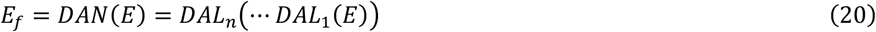

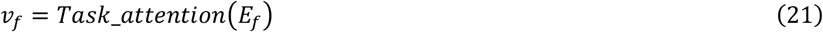

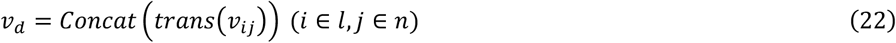

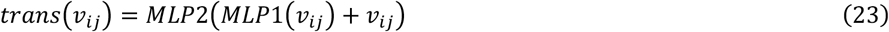

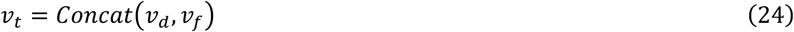

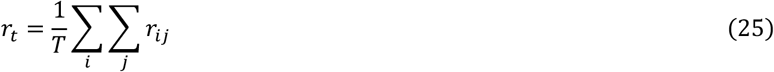

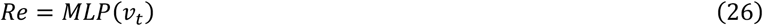

where *E* is the protein representation, *DAL*_*i*_ means the *i*-th Deep Attention Layer, *Task_attention* means the task-specific attention, *v*_*ij*_ is the classification vector generated by *j*-th BAU of *i*-th layer, *trans()* projects *v*_*ij*_ to a vector with a smaller dimension, *l* indicates the number of total attention layers, *n* means the number of BAUs in each attention layer, *Concat* refers to the concatenate function, *v*_*d*_ is the concatenated vector of all classification vectors of BAUs, *E*_*f*_ refers to the output of *DAN, v*_*t*_ is the final classification vector for the model to make prediction, *r*_*t*_ is the integration attention distribution map, *T* is the total number of BAUs in DAN, Re is the prediction result, *MLP,MLP1*, and *MLP2* means Multilayer Perceptron layer.

### Key Segment Generation Algorithm

The deep attention network is utilized to progressively extract attention distribution map across the sequence. The attention at each layer is then integrated by averaging, resulting in a comprehensive attention distribution map *r*_*t*_. This attention distribution map encompasses the attention employed by each unit from various layers. Amino acids in specific positions are sorted according to their attention weights in *r*_*t*_. In our framework, we first define “exploration cofactor” as a parameter for recommendation. This parameter determines the proportion of the full length of a sequence that will be extracted to generate key segments. Our model identifies amino acids that rank in the top proportion of the exploration cofactor in *r*_*t*_. We refer to the collection of extracted amino acids at various positions as pre-selected key positions.

Within the pre-selected set of key positions, an exploration is conducted based on the distances among the key positions to merge amino acids to segments. The criterion for merging two positions is determined based on the distance parameter *H*. Based on the fact that the least length of a NLS is 3, we set *H* to be 2 such that key sites are located at the end and start of a 3-length NLS, it precisely includes the NLS without introducing other non-relevant amino acid. This process gets a collection of important segments by merging pre-selected key positions. These important segments are considered crucial for the DAN’s final prediction, and we refer to them as key segments.

The principle of our algorithm is to find key positions with high attention weights, it cannot guarantee that the key site is the start or end of NLS or another important segment. Simply connecting two key amino acids may lead to inaccurate results. To address this, we adopt a strategy to stretch the segment in a predefined way at the start and end positions. We assume that if a position has a high attention weight, the area surrounding it is also likely to be a key area.

When using a low exploration cofactor, some single residues may not have a neighboring point close enough to form a segment. This issue gradually disappears as the cofactor increase. We also utilize a stretch strategy to solve it. We introduce a sampling policy when taking the stretch strategy and choosing not to filter the single residue. When a key site is located in the middle of an NLS, the stretch length should be longer compared to when it is at the ends of an NLS. In addition, the key segment generation algorithm is based on a predefined template that indicates which BAUs in A2KA contribute to the attention distribution. In this study, we find that using BAUs that prefer to segments enriched with K/R shows better performance than using all the BAUs. The selective mechanism can further help detect specific areas when a suitable template is chosen. We provide the details of the Key Segment Generation Algorithm and the Selective Key Segment Generation Algorithm in the Supplementary File.

### Recommendation system for NLS screening

Upon receiving the key segment set, we further utilize a recommendation system to identify potential NLS segments. Recommendation system of the NLSExplorer is divided into two types. The NLSExplorer-Accuracy adds an initial Mamba^50^ layer to first process the entire protein embedding from ESM, whereas the NLSExplorer-Lightning does not add the Mamba layer. The remaining identical components of both models are consists of a 2-layer Bidirectional Long Short-term Memory (BiLSTM)^51^, a 12-layers Mamba and an Att_mlp layer. BiLSTM can handle the sequence characteristic from two directions. Mamba is good at handling long sequence. In the NLS recommendation task, some NLS segments are long, which makes Mamba a good choice for this scenario.

For a key segment set, we first use ESM-1b to generated the protein representation, and extract the representation for each segment. These representations of segments are fed into our recommendation system to determine the probability of being NLSs.

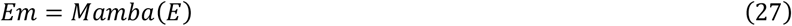

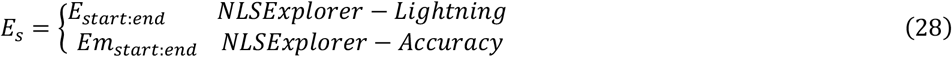

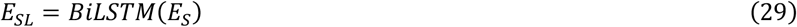

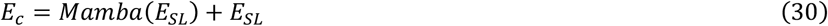

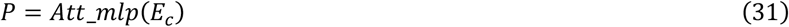

where *E* is the protein representation, *E*_*m*_ indicates the protein representation processed by the initial Mamba layer of NLSExplorer-Accuracy, *E*_*S*_ refers to the extracted segment representation, *BiLSTM* refers to the Bidirectional Long Short-term Memory network, *E*_*SL*_ is the representation processed by *BiLSTM, E*_*c*_ is the segment representation processed by *Mamba, Att_mlp* is an attention classifier network, *P* is the probability of being NLS.

### NLSExplorer Training Details

The model training is divided to two main parts, as shown in Figure 6-c, the first part involves training A2KA on the NLSExplorer-p datasets, A2KA is trained in a supervised way to accurately predict whether a protein locates in nucleus. This ensures that the attention mechanism in A2KA focuses on protein segments that might facilitate nuclear entry. We select A2KA with 16 layers, each layer consists of 64 BAUs. We set the learning rate of 0.00001, batch size of 24, and Adam as the optimizer. The model is trained for 25 epochs on a Titan-X 12GB GPU. During model training, we use all 14,316 positive samples located in the nucleus and 528,826 non-nuclear proteins as negative sample set. For every 5 epochs, we sample 14,316 negative samples from the negative sample set with all the 14,316 positive samples to facilitate the model to see more negative patterns.

The second part involves training the recommendation system in NLSExplorer to further accurately identify NLSs. First, we use A2KA to generate key segment set for the training of the recommendation system. Next, each segment is labeled as a positive sample if it overlaps with a true NLS, otherwise, it is labeled as a negative sample.

We performed 5-fold cross-validation on the training datasets. After 5-fold cross-validation, we trained the model on the INSP training sets and tested its performance on the hybrid and yeast test sets. Finally, we trained our final recommendation system using all NLS data from the INSP datasets and Swiss-Prot with experimental evidence.

### SCNLS (Search and Collect NLS) Pattern Analysis Algorithm

For a given sequence *s*, the objective is to mine implicit patterns within the sequence. For NLS, there exists not only typical contiguous patterns of the Monopartite (MP) type, but also an extensive array of Bipartite (BP) patterns. The NLS of the BP type comprises two non-contiguous segments of core amino acids with a positive charge on both ends. These two positively charged amino acid segments may be separated by a gap, such as “R/K(∗)_10−12_KR*K” (* means any amino acid) with 10-12 amino acids. The amino acids within this gap act as connecting segments, and both non-contiguous amino acid segments must coexist for the NLS to effectively guide the protein into the cell nucleus. Hence, the exploration of non-contiguous patterns holds significant importance in NLS pattern mining. Consequently, a pattern mining algorithm is proposed to identify and explore patterns in both contiguous and non-contiguous NLS, including the potential discovery of NLS types beyond the traditional MP and BP patterns, such as the PYNLS signals formed by N-terminal hydrophobic and C-terminal R/K/H(∗)_2−5_PY motifs ((∗)_2−5_of 2–5 residues).

For a sequence *s* with a length of *l*, we predefine a threshold *M*. We seek all positive integers smaller than or equal to *M*, whose sum equals to *l* forming a set *G*—a mathematical problem known as finding composition of an integer^52^, which can be transformed to inserting blank spaces among *l* balls arranged in a line. For instance, when considering the total compositions of 4, we have: 4, 3 + 1, 1 + 3, 2 + 2, 2 + 1 + 1, 1 + 2 + 1, 1 + 1 + 2, 1 + 1 + 1 + 1. When dealing with a sequence of length 4, there are 8 possible compositions. For example, if we take a hypothetical sequence “MKRK” with a length of 4, based on its composition, it can be partitioned as follows: “MKRK”, “MKR|K”, “M|KRK”, “MK|RK”, “MK|R|K”, “M|KR|K”, “M|K|RK”, “M|K|R|K”. “|” means the separate icon (“blank”) at this point, all possible composition of this NLS is enumerated. Here we only need to determine the positions where the gaps occur and the numbers of gaps, and then we can get all the discontinuous pattern for the target NLS. For instance, given the composition [1, 2, 1], when the gap occurs in the second position, the 3-order length composition becomes [1, *gap*2, 1]. When using this composition on the sequence, we obtain a non-contiguous pattern “M**K”.

For each combination, we derive its subset according to the left-to-right order subset with excluding any repeated subsets. We propose the following approach to identify NLS patterns within NLS. First, we define left-to-*k*-th order subset *G*_*k*_ for a sequence *s* as:

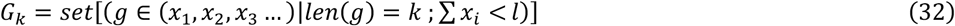

where *g* indicates the composition in *G*_*k*_, *len* () refers to the length of the set, *x*_*i*_ represents the element in *g, l* means the length of *s*.

The computational burden can be alleviated, as each subset needs a single calculation, and they can be recurrently applied to various NLSs, thereby optimizing the efficiency in the analysis. Then, we generate the discontinuous pattern of *G*_*k*_ by applying the gap at various positions:

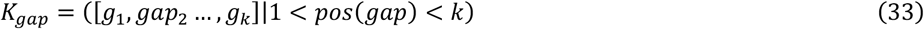

where *K*_*gap*_ denotes the set of gap combination, *pos* () refers to the position of the gap occurrence.

For an element *g*_*k*_ in *G*_*k*_, the combination *K*_*gapSet*_ in *K*_*gap*_ is applied, we define *g*_*k*_ ∗ *K*_*gapSet*_ as the generated non-contiguous combination for the sequence *s*, and *G*_*k*_ ∗ *K*_*gapSet*_ means applying *K*_*gapSet*_ to all elements in *G*_*k*_.

For instance, given the sequence “MSARRK” and the selection of [1, 3, 2] in *G*_3_ and the gap combination of *K*_[1,*gap*2,3]_, the product [1,3,2] ∗ *K*_[1,*gap*2,3]_ = [1, *gap*3,2] Applying this to the sequence yields the pattern “M***RK”. At this point, the notation [1,3,2] signifies the subset of the composition. Initially, the sequence is transformed into “M|SAR|RK”. Then, *K*_[1,*gap*2,3]_ indicates where the gap appears. *gap*2 denotes the gap separated by blanks, and the discontinuous pattern [1, *gap*3,2] alters the sequence to “M|***|RK”. At last, all “blanks” are removed, resulting in “M***RK”.

The BP NLS pattern implied in a key segment set can be obtained by applying *G*_3_ ∗ *K*_[1,*gap*2,3]_ on it and all of its substrings, since it follows the mode in accordance with *K*_[1,*gap*2,3]_, which means a gap in the middle and segments at the ends. We call our algorithm SCNLS (Figure 6-c). In addition to BP NLS, SCNLS can also highlight other patterns. For example, [1, *gap*2,2, *gap*1,3] is a discontinuous pattern that includes two gaps and three continuous core segments. Applying it to a recommendation segment such as “MAAASRKLP” yield a discontinuous pattern of “M**AS*KLP”.

NLSExplorer-prediction obtains a segment set that not only includes high-quality segments for a specific type but also has sufficient volume by utilizing its high exploration capability on the Swiss-Prot database. This provides a valuable reference for SCNLS to mine frequent patterns. By relying on statistical rules, NLSExplorer-SCNLS is able to identify potential NLS patterns implied in key segment sets.

### Maximum Entropy for NLSs

We utilize the maximum entropy for a given amino acid segment to evaluate the diversity of a given NLS, and we can remove those recurrent amino acids with low entropy like ‘QQQQQ’ or ‘AAAAAA,’. In addition, the distribution of maximum entropy for amino acids in specific motifs or domains can help mine new segment patterns.

For a given amino acid segments *s*, its entropy *E*_*s*_ is defined:

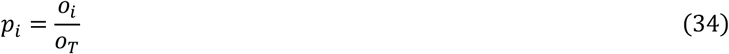

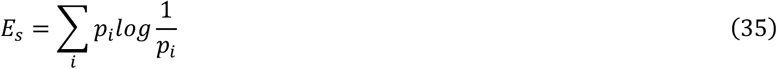

where *p*_*i*_ means the occurrence of amino acid *i, o*_*i*_ means the occurrence of amino acid *i, o*_*T*_ represents the total frequency for all amino acid in the segment *s*.

For NLS, there exist continuous and discontinuous patterns. The continuous pattern can be directly calculated by the formula 35. For discontinuous pattern, we use a method called maximum entropy. The principle of maximum entropy is widely used in various fields like physics^53, 54^, biology^55-57^ and deep learning^58^. In this study, we define the maximum entropy *E*_*ms*_ for a discontinuous pattern as follows:

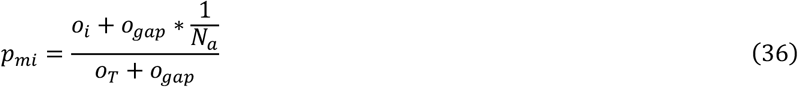

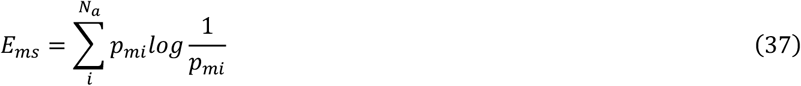

where *o*_*gap*_ means the concurrency of a gap in a discontinuous pattern, *N*_*a*_ is the numbe*r* of amino acids type, *p*_*mi*_ means the occurrence of amino acid *i* under the maximum entropy condition, *o*_*i*_ means the occurrence of amino acid *i, o*_*T*_ represents the total frequency for all amino acid in segment *s*.

For a discontinuous pattern such as “P******RK”, we assume all types of amino acids have an equal probability of occurring in the gap. Then 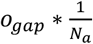 calculates the expectation for all types of amino acid that exists in the gap of the NLS segment and subsequently add to the current amount to derive the final total. Once the occurrence probability for each amino acid is obtained, the entropy of the gap pattern can be derived through the same calculation as Equation 33.

Maximum entropy avoids enumerating all possible combinations for a discontinuous pattern. Although enumerating all combinations and averaging their entropy to get the final entropy may be a more accurate way. It has a high computational complex of O (*N*_*a*_^*ogap*^), especially *o*_*gap*_ is large in the segment. Conversely, the maximum entropy principle provides a more effective way with a linear complex of O (*N*_*a*_) to reliably evaluate entropy.

### Search correlation among species

We use a standard called search correlation to evaluate the NLS correlation among species. First, we generate representations for amino acid segments using a simple TF-IDF (Term Frequency-Inverse Document Frequency). TF-IDF is a standard technique used in sequence analysis that considers both the occurrence and usage frequency of terms. The term frequency highlights frequently occurring segments or amino acids. while the inverse document frequency reduces the total assessment value for amino acids that occur too frequently in each segment. These amino acids are similar to the prepositions function in natural language and cannot convey major information for a segment. TF-IDF has various applications in bioinformatics^59, 60^. Using TF-IDF to represent amino acid segments, we define the following method to calculate the NLS segment correlation among species. For species A and B, we define:

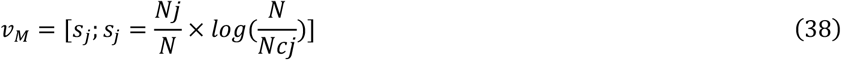

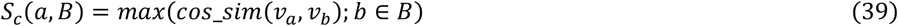

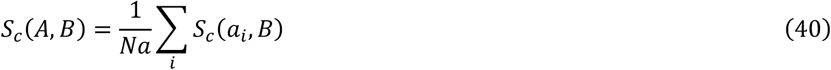

where *v*_*M*_ indicates the representation for amino acid *M, s*_*j*_ represents the value of dimension *j, Nj* is the occurrence of amino acids *j, N* refer to the number of total recommendation segments, *Ncj* indicates the number of recommendation segment that include amino acid *j, cos_sim* is the cosine similarity, *a* and *b* represents amino acid segment, *v*_*a*_ is the representation of amino acid a, *v*_*b*_ is the representation of amino acid *b, A* and *B* indicate the species index, *S*_*c*_(*a, B*) means the search correlation between segment a and species *B, S*_*c*_(*A, B*) indicates the search correlation between species *A* and species *B, Na* is the number of recommendation segments in species *a*.

Due to the limited number of experimentally validated nuclear proteins, some species have insufficient data. As a result, the segment recommendations from these species may not accurately reflect their overall natural appearance. Consequently, it would be inaccurate to place two species in the same positions and calculate the overall coverage rate as their correlation index. However, determining whether other species have similar patterns is feasible and can partially reveal common characteristics. Therefore, we choose search correlation to reflect this. Search correlation implies the extent to which species A can find similar NLS patterns in species B, which means *S*_*c*_(*A, B*) is not necessarily equal to *S*_*c*_(*B, A*).

### Potential NLS visualization

#### NLS candidate library

To further explore the NLS space, we focus on proteins located in the nucleus in Swiss-Prot. We use A2KA in NLSExplorer with an exploration cofactor of 0.3, without filtering single residue or applying stretch strategy, to get key segment sets. We then used the recommendation system of NLSExplorer to assess the probability of each key segment generated by A2KA being an NLS.

We use UMAP to map the representation of the segment into 2D space and employ Plotly and Dash in Python to build an interactive visualization. For the segments obtained from the recommendation system, we project them based on their representation generated by the

*Task_attetion* of *Att_mlp* in the recommendation system. We then utilize the representations of all potential NLS segments to generate a 2D UMAP projection using the default top 5 segments with a probability higher than 0.7.

#### Nuclear Transport Pattern Map

We run the A2KA module of NLSExplorer in a parameter of without filtering single residue and stretching segments in an exploration cofactor of 0.3. Subsequently, NLSExplorer-SCNLS is utilized to analyze the continuous pattern of potential nuclear important segments including NLSs. For a given segment pattern *Se*, we propose the following approach to determine its projection in the two-dimensional coordinate system, where the x-axis represents the hydrophobicity. Since hydrophobicity amino acids play a key role in NLS. They can be integral to the function and regulation of NLSs, which allow for a wide range of sequences to function as NLSs^61^. Therefore, we believe hydrophobicity can be a useful standard to depict the potential pattern.

We set y-axis to represent the maximum entropy for a given pattern. Maximum entropy is a powerful tool to filter noise segment from the frequent pattern sets explored by NLSExplorer. Hence, the y coordinate can be calculated using Formula 35. Then, we derive x coordinate of the segment as follows:

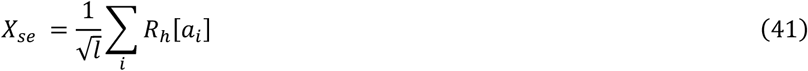

where *X*_*se*_ refers to the x coordinates of the segment pattern *Se, l* refers to the length of *Se, a*_*i*_ refers to the *i*-th amino acid in *Se, R*_*h*_ represent the hydrophobicity value of amino acids.

Following the aforementioned algorithm, we derived the spatial coordinates of the NLS pattern. Subsequently, each point is assigned different sizes and colors based on the corresponding occurrence. Segments pattern that occur frequently exhibit larger point sizes and deeper colors.

#### NLS discontinuous pattern

For a sequence with length *l*, enumerating all discontinuous patterns to discover frequent patterns has a high complexity of O (2^*l*^), compared to continuous patterns with a complexity of O (*l*^2^). This requires an extreme computational resource, especially for mining a large number of sequences. Hence, mining and displaying all discontinuous patterns is impractical. However, it is feasible to gain insights into a specific type of discontinuous pattern in potential mined segments. Thus, we provide the SCNLS codes with multiprocessing capabilities in a Python package with customizable parameters. This approach is efficient when users specify a particular discontinuous pattern to explore.

## Data and code availability

The web-server and source code of NLSExplorer are freely available at www.csbio.sjtu.edu.cn/bioinf/NLSExplorer/ for academic use.

## Acknowledgements

This work is supported by the National Natural Science Foundation of China (No. 62073219, 62473257), the Major Research Plan of the National Natural Science Foundation of China (No. 92059206) and the Science and Technology Commission of Shanghai Municipality (22511104100).

---

**Extended Data Fig 1.**
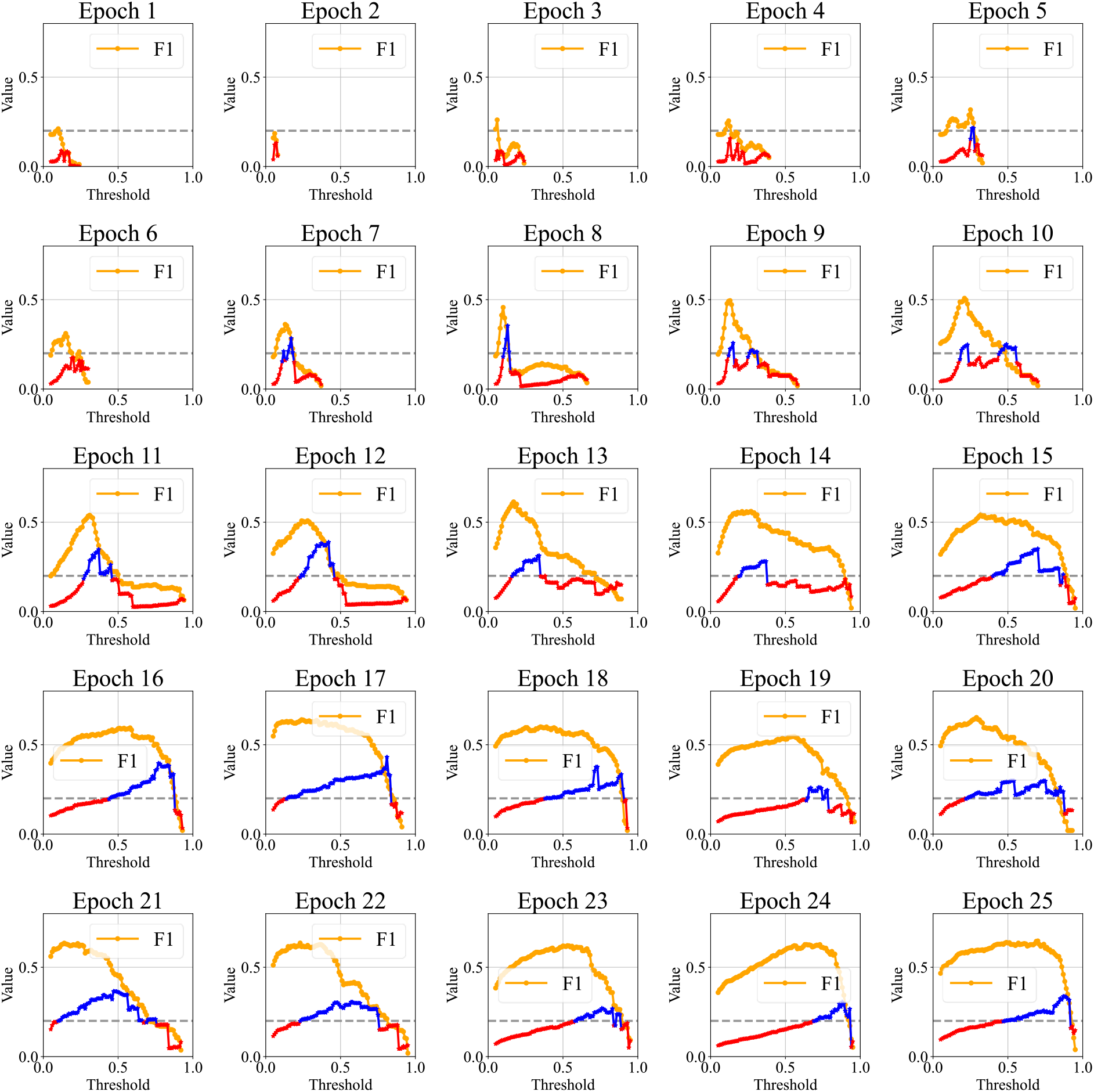
Five-fold cross-validation in INSP training dataset.

**Extended Data Fig. 2.**
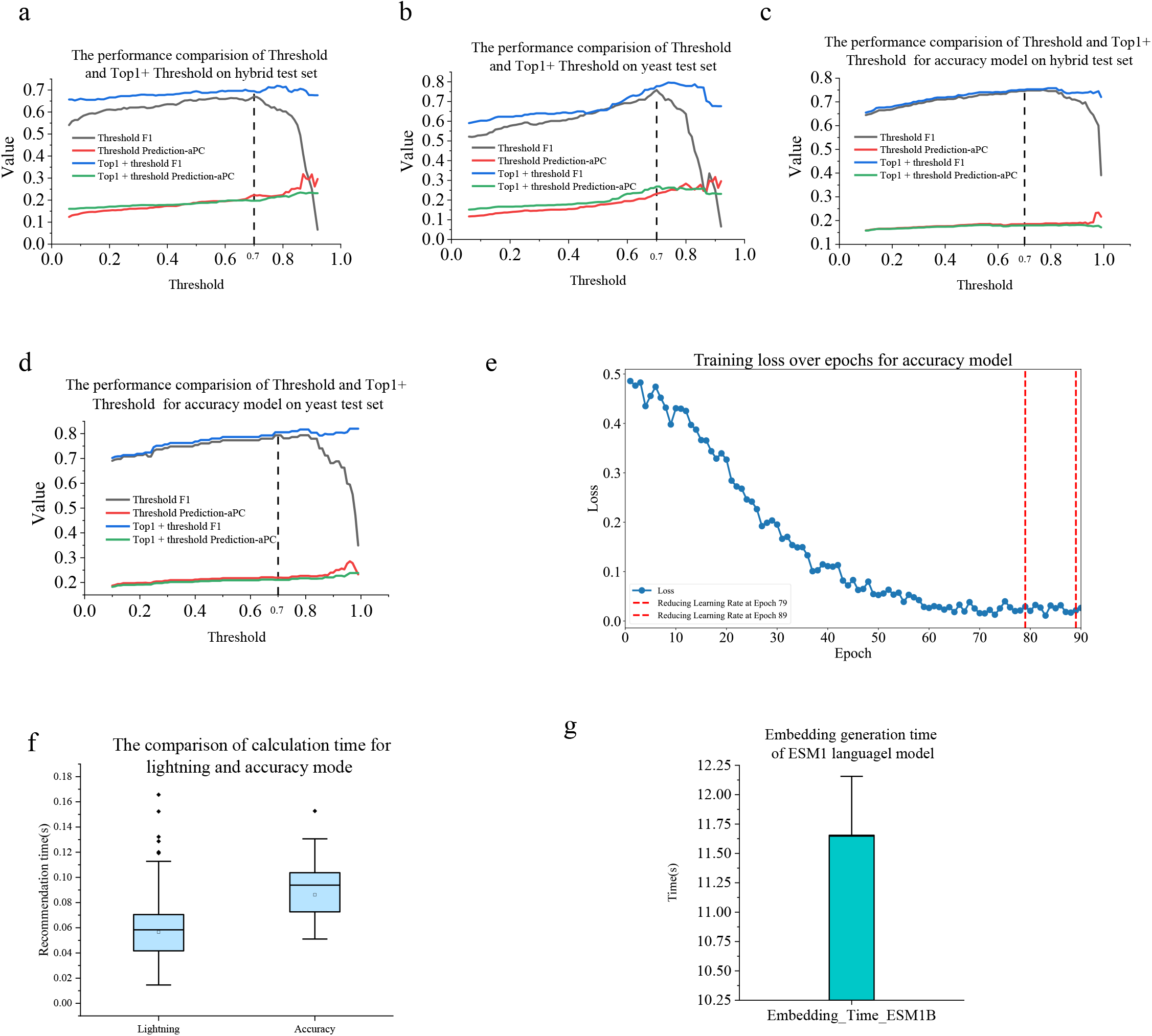
The performance of Top1+Threshold method. **(a)** The F1 and aPC of Top1+Threshold method for NLSExplorer-Lighting on the hybrid test set. **(b)** The F1 and aPC of Top1+Threshold method for the NLSExplorer-Lighting on the yeast test set. **(c)** The F1 and aPC of Top1+Threshold method for the NLSExplorer-Accuracy on the hybrid test set. **(d)** The F1 and aPC of Top1+Threshold method for NLSExplorer-Accuracy on the yeast test set. **(e)** The training process of the NLSExplorer-Accuracy requires a GPU with at least 24GB of memory, compared to 12GB for the NLSExplorer-Lighting, and more epochs to achieve its best performance. We use a dynamic learning rate adjustment strategy and an early stopping mechanism. When the loss over 5 epochs exceeds the previous minimum loss, the learning rate is reduced by multiplying by 0.1. If the learning rate is reduced more than twice, we stop the training. **(f)** The computational time of NLSExplorer-lightning and NLSExplorer-Accuracy when the input sequence length varies from 3 to 1022. **(g)** The embedding generation time of the ESM1 language model using an NVIDIA 3090ti GPU.

**Extended Data Fig. 3.**
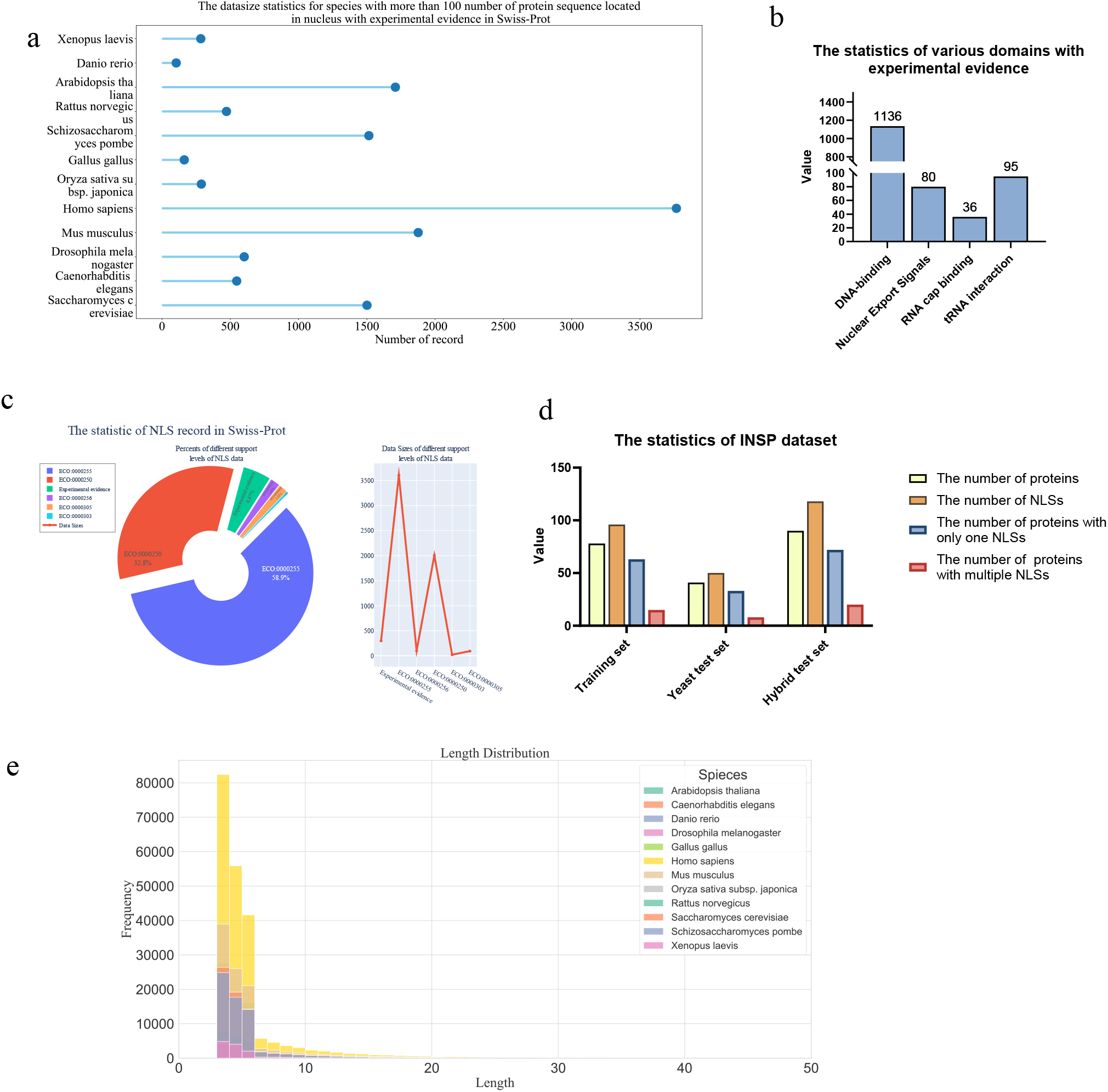
The statistic of the datasets and the length of recommendation segments in this study. **(a)** The data statistics or species with more than 100 protein sequences located in nucleus with experimental evidence in Swiss-Prot. **(b)** The statistic of various domains in Swiss-Prot with experimental evidence. **(c)** The statistic of NLSs in Swiss-Prot database. **(d)** The statistic of INSP dataset. **(e)** The statistic of lengths for recommendation segments generated by NLSExplorer on experimentally validated nuclear proteins in Swiss-Prot. The length of recommendation segments mainly falls in the range of 3 to 50.

**Extended Data Fig. 4.**
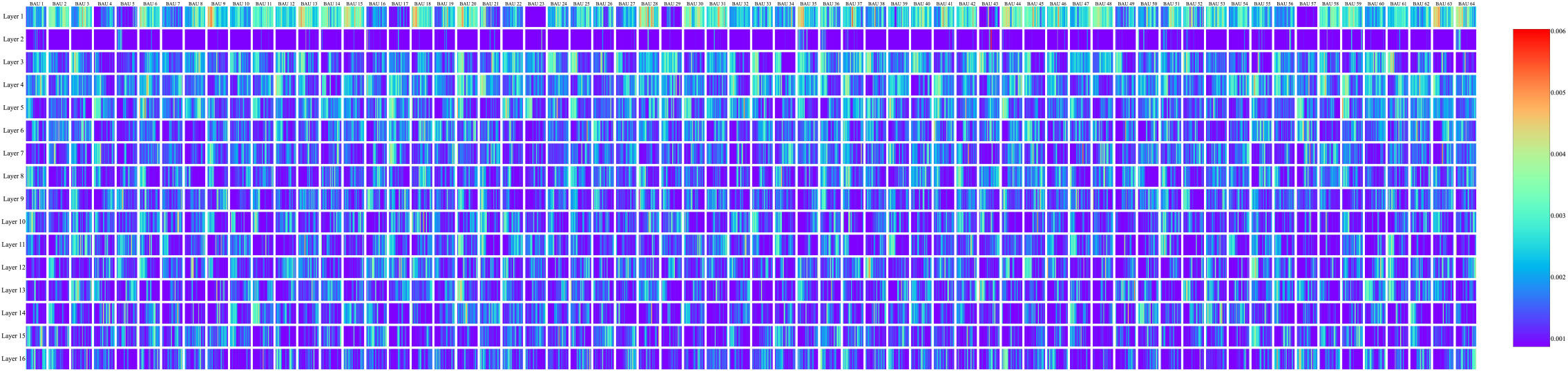
An example of attention map for individual BAUs in DAN on the protein with Swiss-Prot ID Q8N884.

**Extended Data Fig. 5.**
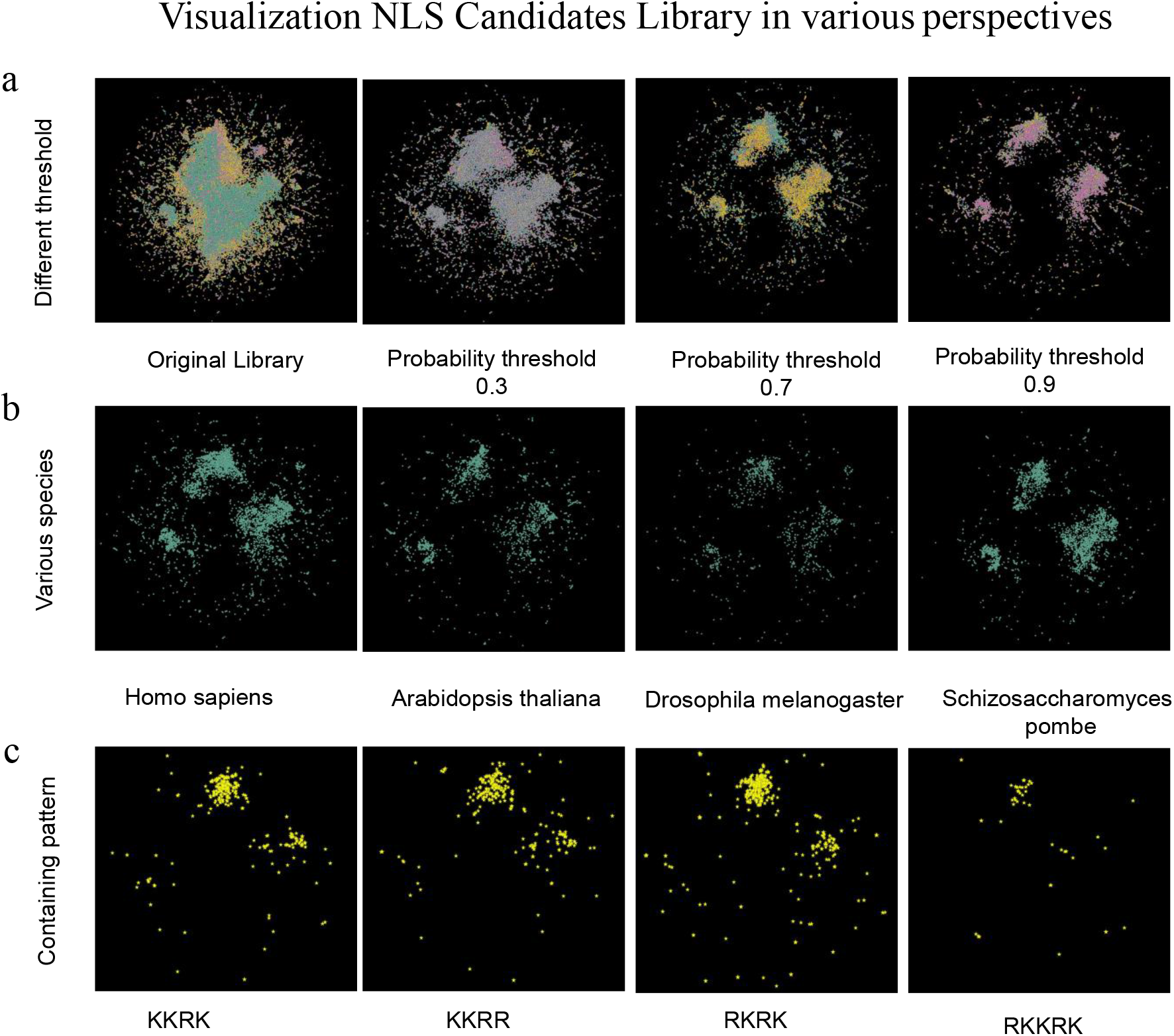
NLS Candidate Library display the potential NLS space from various perspectives. See detail in the chapter “The interactive online application” of Supplementary File. (a) The NLS Candidate Library is constructed using different probability thresholds. The original library is built by running A2KA on all experimentally validated nuclear proteins in Swiss-Prot. The Top 5 segments and segments with a probability higher than 0.7 are selected for visualization. (b) NLS Candidate Library for various species. (c) The NLS Candidate Library displays recommended segments that contain specific amino acid sequences.

## Reference

1. Yu, C.S., Chen, Y.C., Lu, C.H. & Hwang, J.K. Prediction of protein subcellular localization. Proteins: Structure, Function, and Bioinformatics 64, 643–651 (2006).

2. Boulikas, T. Nuclear localization signals (NLS). Critical reviews in eukaryotic gene expression 3, 193–227 (1993).

3. Lu, J. et al. Types of nuclear localization signals and mechanisms of protein import into the nucleus. Cell communication and signaling 19, 1–10 (2021).

4. Yu, M. et al. Visualizing the disordered nuclear transport machinery in situ. Nature, 1-8 (2023).

5. Guang, S. et al. An Argonaute Transports siRNAs from the Cytoplasm to the Nucleus. Science 321, 537–541 (2008).

6. Cai, H. et al. Nucleocytoplasmic Shuttling of a GATA Transcription Factor Functions as a Development Timer. Science 343, 1249531 (2014).

7. Misra, R. & Sahoo, S.K. Intracellular trafficking of nuclear localization signal conjugated nanoparticles for cancer therapy. European Journal of Pharmaceutical Sciences 39, 152–163 (2010).

8. Goswami, R. et al. Nuclear localization signal-tagged systems: relevant nuclear import principles in the context of current therapeutic design. Chemical Society Reviews (2024).

9. Dubrovsky, L. et al. Nuclear localization signal of HIV-1 as a novel target for therapeutic intervention. Molecular Medicine 1, 217–230 (1995).

10. Branden, L.J., Mohamed, A.J. & Smith, C. A peptide nucleic acid-nuclear localization signal fusion that mediates nuclear transport of DNA. Nature biotechnology 17, 784–787 (1999).

11. Salman, H. et al. Nuclear localization signal peptides induce molecular delivery along microtubules. Biophysical journal 89, 2134–2145 (2005).

12. Meier, U.T. & Blobel, G.n. A nuclear localization signal binding protein in the nucleolus. The Journal of cell biology 111, 2235–2245 (1990).

13. Ishidate, T. et al. Identification of a novel nuclear localization signal in Sam68. FEBS letters 409, 237–241 (1997).

14. Xiao, Z., Latek, R. & Lodish, H.F. An extended bipartite nuclear localization signal in Smad4 is required for its nuclear import and transcriptional activity. Oncogene 22, 1057–1069 (2003).

15. Consortium, T.U. UniProt: the Universal Protein Knowledgebase in 2023. Nucleic Acids Research 51, D523–D531 (2022).

16. Vinokourov, A., Cristianini, N. & Shawe-Taylor, J. Inferring a semantic representation of text via cross-language correlation analysis. Advances in neural information processing systems 15 (2002).

17. Alley, E.C., Khimulya, G., Biswas, S., AlQuraishi, M. & Church, G.M. Unified rational protein engineering with sequence-based deep representation learning. Nature methods 16, 1315–1322 (2019).

18. Lin, Z. et al. Evolutionary-scale prediction of atomic-level protein structure with a language model. Science 379, 1123–1130 (2023).

19. Cheng, J. et al. Accurate proteome-wide missense variant effect prediction with AlphaMissense. Science 381, eadg7492 (2023).

20. Teufel, F. et al. SignalP 6.0 predicts all five types of signal peptides using protein language models. Nature biotechnology 40, 1023–1025 (2022).

21. Thumuluri, V., Almagro Armenteros, J.J., Johansen, A.R., Nielsen, H. & Winther, O. DeepLoc 2.0: multi-label subcellular localization prediction using protein language models. Nucleic Acids Res 50, W228–w234 (2022).

22. Vig, J. et al. Bertology meets biology: Interpreting attention in protein language models. arXiv preprint arXiv:2006.15222 (2020).

23. Chowdhary, K. & Chowdhary, K. Natural language processing. Fundamentals of artificial intelligence, 603-649 (2020).

24. Rives, A. et al. Biological structure and function emerge from scaling unsupervised learning to 250 million protein sequences. Proceedings of the National Academy of Sciences 118, e2016239118 (2021).

25. Suzek, B.E., Huang, H., McGarvey, P., Mazumder, R. & Wu, C.H. UniRef: comprehensive and non-redundant UniProt reference clusters. Bioinformatics 23, 1282–1288 (2007).

26. Hodel, M.R., Corbett, A.H. & Hodel, A.E. Dissection of a nuclear localization signal. Journal of Biological Chemistry 276, 1317–1325 (2001).

27. Guo, Y., Yang, Y., Huang, Y. & Shen, H.-B. Discovering nuclear targeting signal sequence through protein language learning and multivariate analysis. Analytical biochemistry 591, 113565 (2020).

28. Kosugi, S., Hasebe, M., Tomita, M. & Yanagawa, H. Systematic identification of cell cycle-dependent yeast nucleocytoplasmic shuttling proteins by prediction of composite motifs. Proc Natl Acad Sci U S A 106, 10171–10176 (2009).

29. Schäfer, F., Florin, L. & Sapp, M. DNA binding of L1 is required for human papillomavirus morphogenesis in vivo. Virology 295, 172–181 (2002).

30. Prieve, M.G., Guttridge, K.L., Munguia, J. & Waterman, M.L. Differential importin-alpha recognition and nuclear transport by nuclear localization signals within the high-mobility-group DNA binding domains of lymphoid enhancer factor 1 and T-cell factor 1. Mol Cell Biol 18, 4819–4832 (1998).

31. Fukao, A., Tomohiro, T. & Fujiwara, T. Translation Initiation Regulated by RNA-Binding Protein in Mammals: The Modulation of Translation Initiation Complex by Trans-Acting Factors. Cells 10, 1711 (2021).

32. Taggart, D.J., Mitchell, J.K. & Friesen, P.D. A conserved N-terminal domain mediates required DNA replication activities and phosphorylation of the transcriptional activator IE1 of Autographa californica multicapsid nucleopolyhedrovirus. J Virol 86, 6575–6585 (2012).

33. Jumper, J. et al. Highly accurate protein structure prediction with AlphaFold. Nature 596, 583–589 (2021).

34. Xu, J. & Zhang, Y. How significant is a protein structure similarity with TM-score = 0.5? Bioinformatics 26, 889–895 (2010).

35. Kee, K., Angeles, V.T., Flores, M., Nguyen, H.N. & Reijo Pera, R.A. Human DAZL, DAZ and BOULE genes modulate primordial germ-cell and haploid gamete formation. Nature 462, 222–225 (2009).

36. Liu, H. et al. Nuclear cGAS suppresses DNA repair and promotes tumorigenesis. Nature 563, 131–136 (2018).

37. Wang, Y. et al. Inflammasome Activation Triggers Caspase-1-Mediated Cleavage of cGAS to Regulate Responses to DNA Virus Infection. Immunity 46, 393–404 (2017).

38. Yang, Y. et al. WW domains form a folded type of nuclear localization signal to guide YAP1 nuclear import. Journal of Cell Biology 223 (2024).

39. Lin, J.-r. & Hu, J. SeqNLS: nuclear localization signal prediction based on frequent pattern mining and linear motif scoring. PloS one 8, e76864 (2013).

40. Chang, Y. et al. A survey on evaluation of large language models. ACM Transactions on Intelligent Systems and Technology (2023).

41. Guo, Y., Wu, J., Ma, H. & Huang, J. in Proceedings of the AAAI Conference on Artificial Intelligence, Vol. 36 6801–6809 (2022).

42. Greener, J.G. & Jamali, K. Fast protein structure searching using structure graph embeddings. bioRxiv, 2022.2011. 2028.518224 (2022).

43. Pan, X. et al. ToxDL: deep learning using primary structure and domain embeddings for assessing protein toxicity. Bioinformatics 36, 5159–5168 (2020).

44. van Kempen, M. et al. Fast and accurate protein structure search with Foldseek. Nature Biotechnology 42, 243–246 (2024).

45. Xia, C., Feng, S.H., Xia, Y., Pan, X. & Shen, H.B. Fast protein structure comparison through effective representation learning with contrastive graph neural networks. Plos Comput Biol 18, e1009986 (2022).

46. Bernhofer, M. et al. NLSdb-major update for database of nuclear localization signals and nuclear export signals. Nucleic acids research 46, D503–D508 (2018).

47. Taud, H. & Mas, J. Multilayer perceptron (MLP). Geomatic approaches for modeling land change scenarios, 451-455 (2018).

48. Rao, R.M. et al. in Proceedings of the 38th International Conference on Machine Learning, Vol. 139. (eds. M. Marina & Z. Tong) 8844-8856 (PMLR, Proceedings of Machine Learning Research; 2021).

49. He, K., Zhang, X., Ren, S. & Sun, J. in 2016 IEEE Conference on Computer Vision and Pattern Recognition (CVPR) 770–778 (2016).

50. Gu, A. & Dao, T. Mamba: Linear-time sequence modeling with selective state spaces. arXiv preprint arXiv:2312.00752 (2023).

51. Zhou, P. et al. in Proceedings of the 54th annual meeting of the association for computational linguistics (volume 2: Short papers) 207–212 (2016).

52. Andrews, G.E. & Eriksson, K. Integer partitions. (Cambridge University Press, 2004).

53. Vartiainen, E.M. & Peiponen, K.E. Optical and terahertz spectra analysis by the maximum entropy method. Rep Prog Phys 76, 066401 (2013).

54. Ezaki, T., Watanabe, T., Ohzeki, M. & Masuda, N. Energy landscape analysis of neuroimaging data. Philos Trans A Math Phys Eng Sci 375 (2017).

55. Stein, R.R., Marks, D.S. & Sander, C. Inferring Pairwise Interactions from Biological Data Using Maximum-Entropy Probability Models. PLoS Comput Biol 11, e1004182 (2015).

56. Esposito, R., Mensitieri, G. & de Nicola, S. Improved maximum entropy method for the analysis of fluorescence spectroscopy data: evaluating zero-time shift and assessing its effect on the determination of fluorescence lifetimes. Analyst 140, 8138–8147 (2015).

57. Weistuch, C., Zhu, J., Deasy, J.O. & Tannenbaum, A.R. The maximum entropy principle for compositional data. BMC Bioinformatics 23, 449 (2022).

58. He, Z., Liu, J., Dang, K., Zhuang, F. & Huang, Y. Leveraging maximum entropy and correlation on latent factors for learning representations. Neural Netw 131, 312–323 (2020).

59. Ranjan, A., Fernandez-Baca, D., Tripathi, S. & Deepak, A. An Ensemble Tf-Idf Based Approach to Protein Function Prediction via Sequence Segmentation. IEEE/ACM Transactions on Computational Biology and Bioinformatics 19, 2685–2696 (2022).

60. Caldas, J. & Vinga, S. Global meta-analysis of transcriptomics studies. PLoS One 9, e89318 (2014).

61. Kosugi, S. et al. Six Classes of Nuclear Localization Signals Specific to Different Binding Grooves of Importin α. Journal of Biological Chemistry 284, 478–485 (2009).

